# The structure of the lipid II flippase from monoderm bacteria

**DOI:** 10.64898/2026.06.21.733627

**Authors:** Yancheng E. Li, Grace F. Baron, William M. Clemons

## Abstract

Peptidoglycan biogenesis requires membrane flippases to translocate lipid-linked precursors across the cytoplasmic membrane for processing (1). This essential step is mediated by MurJ, the lipid II flippase conserved across all peptidoglycan-producing bacteria (2). While MurJ from diderm bacteria has been structurally resolved in multiple conformational states (3–6), its monoderm homolog remains uncharacterized. Monoderm MurJ homologs exhibit substantial sequence divergence yet retain the same lipid II flipping function (7) and are promising antibiotic targets. Here we report structures of *Staphylococcus aureus* MurJ (*Sa*MurJ) captured in both outward- and inward-facing conformations. These structures show that *Sa*MurJ adopts the conserved MOP family fold and undergoes conformational transitions consistent with an alternating-access mechanism. Our findings reveal conserved and divergent features of MurJ between diderm and monoderm bacteria that are critical for lipid II flipping and provide a structural framework for probing substrate recognition and specific inhibition.

**Significance Statement:** The growing global threat of antibiotic resistance and the limited development of new antibacterial therapies underscore the urgent need to identify and mechanistically characterize new antibiotic targets and mechanisms. MurJ is an essential membrane transporter required for cell wall biosynthesis and represents an attractive but unexplored antibiotic target. Here we determine the structures of MurJ from a clinically critical monoderm pathogen *Staphylococcus aureus* in key conformational states during its transport cycle. This work advances our understanding of an essential step in bacterial cell wall synthesis, reveals key distinctions between monoderm and diderm MurJ, and defines structural features that can be exploited for antibiotic discovery.

## Introduction

The biosynthesis of many essential cellular glycoconjugates depends on the translocation of lipid-linked glycan precursors across biological membranes. These lipid-linked intermediates are critical for the biogenesis of diverse glycans, including peptidoglycan, N-linked glycans, and O-antigens (8, 9). The transport step across the membrane bilayer is mediated by lipid flippases (10). In bacterial cell wall biosynthesis, the key precursor to peptidoglycan (PG) biogenesis is lipid II (1), a 55-carbon polyisoprenol unit attached via a diphosphate to a peptide-disaccharide, often a MurNAc-pentapeptide-GlcNAc. Lipid II is made in the cytoplasm, but its head group must be translocated across the cytoplasmic membrane where it is incorporated into PG. The flippase for the lipid II headgroup was first identified in diderms as the conserved and essential MurJ (2). Historically, the search for the equivalent MurJ activity in monoderm bacteria was difficult, as sequence-based searches identified only low-homology MurJ candidates, such as YtgP, that were not essential in *B. subtilis* (11), although evidence supported that YtgP was the MurJ homolog as the gene from *Streptococcus pyogenes* could rescue an *E. coli* MurJ conditional knock-out (7). Definitive evidence that monoderm YtgP was the MurJ homolog came with the discovery that a unique *Bacillus* gene, *amj,* was a redundant lipid II flippase (12). These findings highlight that lipid-linked precursor flippases can remain functionally conserved despite substantial sequence divergence, making them challenging to identify through sequence homology, as exemplified by the long-standing uncertainty surrounding the Rft1 flippase in eukaryotic N-linked glycosylation (13).

MurJ, an integral membrane protein, contains 14 transmembrane helices (TM) and belongs to the MOP (<u>m</u>ultidrug/<u>o</u>ligosaccharidyl-lipid/<u>p</u>olysaccharide) transporter superfamily (3–5). Multiple conformational states of the thermophilic *Thermosipho africanus* MurJ (*Ta*MurJ) have been structurally characterized (5) (Fig. S1), capturing snapshots along the inward-open to outward-open transitions supporting an alternating-access model. The first 12 TMs form the conserved MOP transporter core and are organized into pseudo-C2 symmetric six-helix N- and C-terminal lobes (Fig. S1A). The last two TMs, TM13 and TM14, are unique to MurJ homologs and form a putative binding groove for the polyisoprenyl chain of lipid II (3–5). The model posits that the lipid extends through a lateral membrane portal between TM1 and TM8 to the central cavity, where a conserved and essential arginine triad coordinates the diphosphate moiety of the headgroup (3, 14, 15). In the alternating-access model, lipid II in the cytoplasmic leaflet is captured by the inward-open conformation (Fig. S1, B and C). Substrate recognition in the central cavity, involving the conserved catalytic triad, is thought to promote conformational changes (16, 17). These rearrangements are accompanied by the closure of the membrane portal between TM1 and TM8 (Fig. S1D) and the formation of a thin gate between the G/A-E-G-A motif on TM2, specifically conserved glutamate residue, and R352 (in *Ta*MurJ), which transitions from an open to an occluded state to restrict access to the central cavity (Fig. S1C). This is coupled to sealing of the cytoplasmic gate while opening the periplasmic gate, producing the outward-facing state (5) (Fig. S1B). This exposes the central cavity to the periplasm, allowing lipid II release and subsequent incorporation into peptidoglycan. Similarly, in *E. coli,* MurJ has been captured in an inward-facing (4) and a squeezed conformation (6) (Fig. S1E), although the role of the squeezed state in the transport cycle remains unclear.

MurJ has emerged as a major target for antibiotic discovery (18–21). Multiple screening campaigns have identified compounds active against monoderm pathogens such as *Staphylococcus aureus* and *Streptococcus pneumoniae*, with genetic evidence strongly supporting MurJ as the cellular target (18–21). These inhibitors include small molecules from large synthetic chemical libraries (18, 19) as well as natural products such as the humimycins (20, 21). Moreover, our recent work reveals that MurJ is inhibited by convergently evolved bacteriophages through a common mechanism (22), further highlighting MurJ as an evolutionarily validated key target in bacterial cell wall biosynthesis. While diderm MurJ has been structurally resolved in multiple conformational states (3–6), the monoderm MurJ homolog YtgP remains structurally uncharacterized. Determining the structure of monoderm MurJ is critical for understanding how it diverges from the diderm MurJ and for elucidating the mode of action of inhibitors that specifically target monoderm MurJ.

Here we present the first structures of the MurJ homolog from *S. aureus* (SAV1754, referred to here as *Sa*MurJ) determined by cryogenic electron microscopy (cryo-EM) in both outward- and inward-facing states. We also obtained the structure of an unliganded *E. coli* MurJ in an outward-facing state, enabling direct comparison of MurJ from pathogenic monoderm and diderm bacteria across conformational states. We computationally identified conserved features of the monoderm MurJ lipid II-binding pocket and experimentally probed the residues in the substrate-binding sites. Our findings address a longstanding question regarding conservation of MurJ in monoderm bacteria and establish a structural framework for the development of monoderm- and diderm-selective MurJ-targeting antibiotics.

## Results

### Structure of *Sa*MurJ in an outward-facing state

We first sought to determine the structure of *Sa*MurJ using cryo-EM, whose size (∼55 kDa) makes it a challenging target. We previously overcame this by using a four-helix bundle fusion strategy (23) to determine the structures of *E. coli* MurJ bound to phage-encoded inhibitors (22). While a *Sa*MurJ version of this construct could be purified, we were unable to obtain a high-resolution reconstruction, possibly due to the flexibility of the fusion tag. As an alternative, we used the ALFA-tag system (24), appending the ALFA peptide to the C-terminus of *Sa*MurJ (*Sa*MurJ_ALFA_) (Fig. S2A) and forming a complex with an anti-ALFA-tag nanobody (24) (Fig. S2B).

We determined the structure of *Sa*MurJ_ALFA_ to 3.6 Å using the nanobody as a fiducial (Fig 1A, Fig. S3). Despite low sequence conservation between diderm and monoderm MurJ (17% identity between *Sa* and *Ec*), the structure showed that *Sa*MurJ has a canonical MOP transporter core consisting of 12 TMs in two lobes (N- TM1-6 and C TM 7-12) as well as the two additional helices TM13-14 (Fig. 1, B and C). For this construct, *Sa*MurJ_ALFA_ adopts an <u>o</u>utward-<u>f</u>acing <u>s</u>tate (OFS) with the central cavity formed by TM1, TM2, TM7, and TM8 exposed to the extracellular side (Fig. 1D). TM13 and TM14 form the groove that has been proposed to bind the polyisoprenol chain of the lipid II substrate. The structure demonstrates conservation of the MurJ architecture across bacteria.

**Fig. 1.**
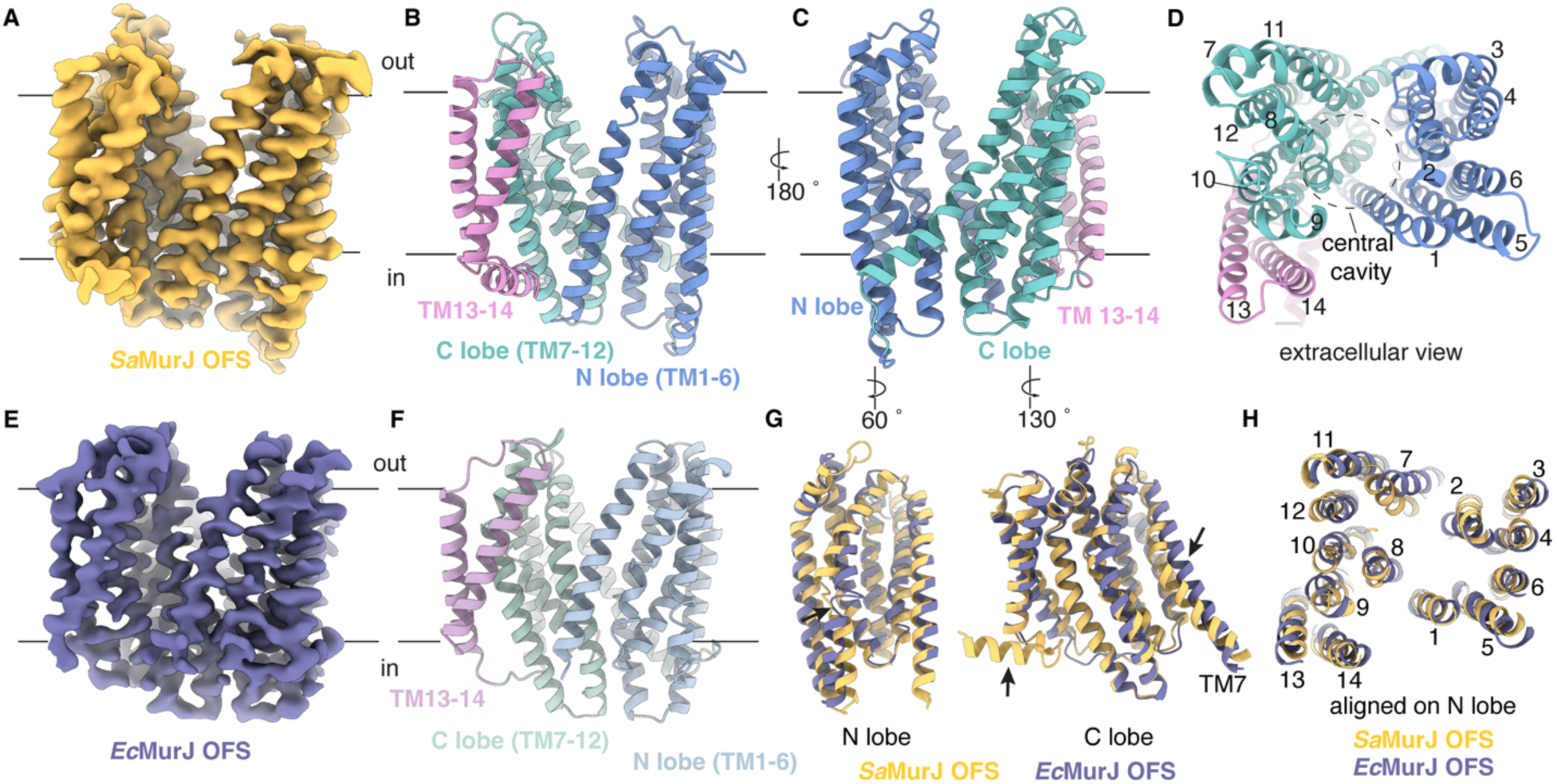
The structure of *Sa*MurJ and *Ec*MurJ in an outward-facing state. (**A**) Cryo-EM map (sharpened) of ALFA-tagged *Sa*MurJ. Approximate membrane boundaries are represented with black lines in all figure panels. (**B**) Cartoon representation of *Sa*MurJ in an outward-facing state in the same orientation as in (A) and colored by domain architecture, with the N-lobe, C-lobe, and TM13-14 shown in blue, green, and pink, respectively. **(C)** As in (B) rotated 180 °. (**D**) Extracellular view of outward-facing *Sa*MurJ highlighting the central cavity (dashed circle). **(E)** Cryo-EM map (sharpened) of ALFA-tagged *Ec*MurJ. (**F**) Cartoon representation of *Ec*MurJ in an outward-facing state colored by domain architecture. (**G**) Structural overlay of N- and C-lobes of *Sa*MurJ and *Ec*MurJ in the outward-facing conformation, colored as in (A) and (E). **(H)** Structural overlay of the outward-facing *Sa*MurJ and *Ec*MurJ aligned on the N-lobe viewed from the extracellular side.

### Structure of *Ec*MurJ in an outward-facing state

To enable a full comparison across pathogenic MurJ homologs, we determined the structure of apo *Ec*MurJ in an outward-facing conformation to 3.9 Å using the ALFA-tag system (Fig. 1E, Fig. S2, Fig S4). This outward-facing conformation is similar to the previously reported outward-facing *Ta*MurJ (5) and the Sgl-bound *Ec*MurJ (22) structures (Fig. 1F).

This outward-facing structure of *Ec*MurJ enables direct comparison with the previously reported inward-facing *Ec*MurJ conformation (PDB: 6CC4), thereby allowing visualization of the structural rearrangements underlying the alternating-access mechanism (25). Comparing the individual lobes reveals the expected rocker-switch mechanism (26), with an overall rigid-body movement between the N- and C-lobes (Fig. S5A). Within the lobes, distinct differences occur in the pseudo-symmetric pairs TM1 & 2 and TM7 & 8 (Fig. S5A). TM1 adopts a straightened helix in the outward-facing state that closes the lateral membrane portal, whereas it appears distorted in the inward-facing crystal structure, possibly caused by the BRIL fusion (Fig. S5A) (4).

During the inward-to-outward transition, TM2 and TM8 shift toward the central cavity, reshaping the cavity architecture. Meanwhile, TM7 straightens, and the TM6-7 loop connecting the N- and C-lobes shifts downward towards the cytoplasmic side. This movement brings the N- and C-lobes closer at the cytoplasmic side, thereby sealing the cytoplasmic gate and occluding access to the central cavity from the cytoplasm. As a result, the central cavity becomes exposed to the periplasm, allowing substrate release. Similar conformational rearrangements during the inward-to-outward transition have also been observed in *T. africanus* MurJ (5).

A structural alignment of the OFS *Sa*MurJ and *Ec*MurJ reveals they have similar overall architecture (Fig. 1, G and H, Fig. S5B) (RMSD for equivalent residues of 1.345 Å). TMs1, 5, 6, 7 & 8 are longer in *Sa*MurJ compared to *Ec*MurJ (Fig. S4B). A more pronounced divergence is observed in the TM6-7 loop, which in *Sa*MurJ extends substantially farther from the membrane compared to *Ec*MurJ (Fig. S5B). Overlay of the individual N- and C-lobes shows that the overall folds are highly similar (RMSD of 1.15 Å and 1.14 Å for N- and C-lobe, respectively), with differences primarily in the positioning of TM2 and TM7 (Fig. 1, G and H). Specifically, TM2 in *Sa*MurJ is shifted more outward from the central cavity, with a corresponding change in the positioning of TM7, which is closely packed against TM2 (Fig. 1G, Fig. S5C). The conformational difference in TM2 occurs at the position corresponding to the diderm conserved G/A-E-G-A motif in *Ec*MurJ (3, 5, 9), which is absent in *Sa*MurJ (Fig. S5C). In *Ec*MurJ, this motif within TM2 introduces a helix break, and mutations here abolish MurJ function (5). This structural flexibility is thought to facilitate TM2 bending and enable rearrangements of TM2 as MurJ switches between inward- and outward-facing states (5). Since *Sa*MurJ lacks the canonical G/A-E-G-A motif, we speculate that *Sa*MurJ might utilize alternative structural features in TM2 to accommodate similar conformational changes. In particular, the highly conserved residues G62 and P64 break the TM2 helix, which likely contributes to the local structural flexibility required for TM2 bending.

### Structure of *Sa*MurJ in an inward-facing state

We next aimed to capture *Sa*MurJ in an inward-facing state to better understand the conformational transitions during the lipid II transport cycle. As initial attempts with full-length *Sa*MurJ fused to BRIL did not yield high-resolution reconstructions, likely due to flexibility, we deleted the first seven N-terminal residues of *Sa*MurJ (*Sa*MurJ^8-553^_BRIL_), which resulted in a more rigid attachment of BRIL to TM1 of *Sa*MurJ^8-553^ and enabled a high-resolution reconstruction to 3.5 Å (Fig. 2A, fig. S6). This construct was in an inward-facing state (IFS) with the central cavity open to the cytoplasm (Fig. 2B).

**Fig. 2.**
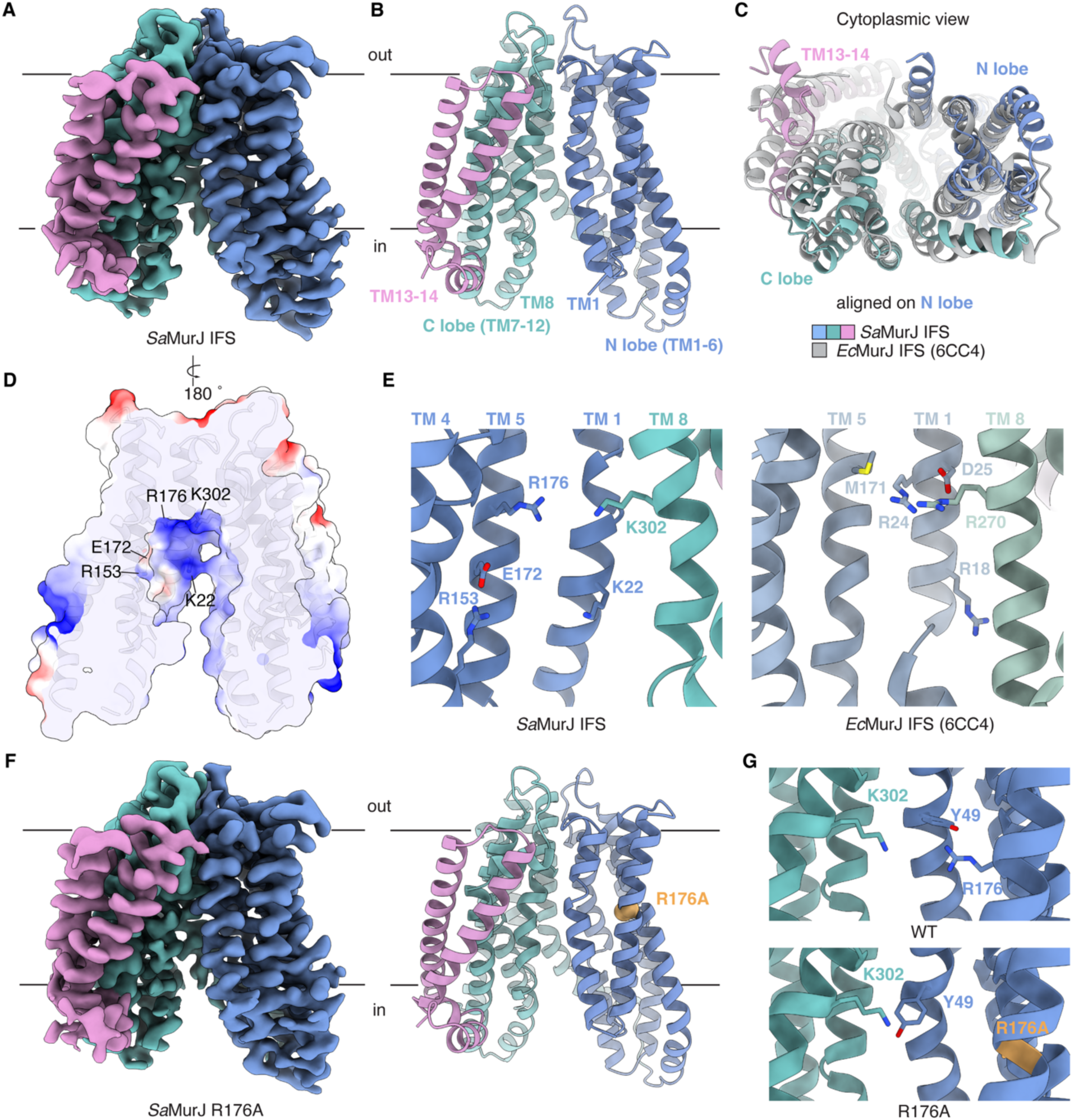
The structure of *Sa*MurJ in an inward-facing state. (**A**) Cryo-EM map (sharpened) of BRIL-tagged *Sa*MurJ_8-553_ colored by domain architecture as in Fig. 1. (**B**) Cartoon representation of *Sa*MurJ_8-553_ in an inward-facing state, colored as in (A). (**C**) Structural overlay of *Sa*MurJ and *Ec*MurJ (PDB:6CC4) in the inward-facing conformation viewed from the cytoplasm. The two structures are aligned on the N-lobe. (**D**) Electrostatic potential surface representation of the central cavity, colored from negative (red) to positive (blue), ranging from -10 to +10 kcal/(mol·*e*). Functionally important charged residues in the central cavity are indicated by lines and labeled. (**E**) Left, close-up view of the charged residues in *Sa*MurJ highlighted in (D), with side chains shown in stick representation. Right, residues in *Ec*MurJ at the corresponding positions. (**F**) Cryo-EM map (sharpened) and cartoon representation of *Sa*MurJ_ALFA_ R176A mutant colored as in (A). R176A is shown in orange. (**G**) Close-up view of the R176A mutant (bottom) compared with WT (top), highlighting residues K302 and Y49.

In the inward-facing state, *Sa*MurJ and *Ec*MurJ adopt similar overall conformations, with some differences in the relative orientation of their N- and C-lobes (Fig. 2C). Unlike *Ec*MurJ, TM1 in *Sa*MurJ is not distorted by the BRIL fusion and instead forms a partially open membrane portal between TM1 and TM8 (Fig. 2C). Consistent with the outward-facing structures, the N- and C-lobes of *Sa*MurJ and *Ec*MurJ overlay well in the inward state, with the same differences observed in the positioning of TM2 and TM7 (Fig. S5D). The functional importance of charged residues in the central cavity of *Ec*MurJ and *Streptococcus pyogenes* MurJ has been tested to show that monoderm and diderm MurJ homologs have unique charge requirements for flippase function (15). Similar to diderm MurJ, *Sa*MurJ has a strongly cationic site (K22, R176, and K302) adjacent to the membrane portal (Fig. 2D), forming a triad of positive residues required for flippase function and proposed to coordinate the diphosphate moiety of lipid II (15). Our structures confirmed that the Lys22-Arg176-Lys302 triad in *Sa*MurJ is structurally equivalent to the Arg18-Arg24-Arg270 triad in *Ec*MurJ (Fig. 2E). R176 in *Sa*MurJ is located at the position equivalent to M171 in *Ec*MurJ, but its side chain extends to a similar position as R24 in *Ec*MurJ, consistent with the ability of an M171R substitution in *E. coli* to suppress the loss-of-function phenotype caused by the R24A substitution (15). Substitution of two additional charged residues in the central cavity of *Sa*MurJ that are proposed to interact with the substrate, in TM4 (R153) and TM5 (E172), resulted in a loss-of-function phenotype (15) (Fig. 2, D and E).

The essential Arg18-Arg24-Arg270 triad in *Ec*MurJ has been proposed to induce inward-to-outward conformational transition in response to substrate recognition within the central cavity (17). We therefore hypothesized that alanine substitutions of these residues would trap MurJ in an inward-facing state. As the ALFA-tagged *Sa*MurJ construct adopts an outward-facing conformation, we introduced an alanine substitution at Arg176 (functionally equivalent to Arg24 in *Ec*MurJ) and determined the structure of *Sa*MurJ_ALFA_ R176A variant to 3.6 Å (Fig 2F, Fig. S7). The *Sa*MurJ_ALFA_ R176A adopts an inward-facing conformation that is similar to the inward-facing structure captured using the BRIL fusion (Fig. 2F). We observed that Y49, conserved in monoderm MurJ, adopts distinct side chain conformations between the wild-type and R176A structures (Fig. 2G). In the wild-type structure, Y49 is positioned to form a cation-π interaction with R176, whereas in the mutant, Y49 shifts away from the R176A (Fig. 2G).

### Conformational transitions of *Sa*MurJ during the lipid II transport cycle

We compared the outward- and inward-facing structures of *Sa*MurJ to assess the conformational transitions that enable MurJ to flip lipid II across the inner membrane. Aligning the C-lobes of the two conformational states reveals that, relatively, the N-lobe undergoes substantial movement, altering the accessibility of the central cavity between the cytoplasm and the extracellular side (Fig. 3A). These conformational rearrangements support a previously proposed model of MurJ-mediated lipid II flipping, in which lipid II is recognized and captured from the cytoplasm in the inward-facing state and translocated through the central cavity via alternating-access conformational transitions before release into the periplasm (5, 9). Alignment of individual lobes shows that the conformational transition arises mainly from relative rigid-body movement between the N- and C-lobes, with the most notable intradomain changes observed in TM1 of the N-lobe and TM7 of the C-lobe (Fig. 3B). TM1 shifts towards the central cavity and closes the lateral membrane portal as *Sa*MurJ transitions to the outward-facing state (Fig. 3B), likely induced in response to substrate binding. In parallel, TM7 undergoes additional rotational movement, lowering the position of the TM6-7 loop relative to the membrane (Fig. 3B). These conformational changes described above are similar to those observed in *Ec*MurJ, highlighting the conservation of the alternating-access mechanism. In both the outward- and inward-facing structures, several salt bridges are observed at the gate regions, where they are positioned to stabilize the corresponding conformational states (Fig. 3, C and D). These include interactions at the cytoplasmic gate between E3-K324 and K69-D411 in the outward-facing structure (Fig. 3C), and at the extracellular gate between R119-D290 in the cytoplasm-open structure (Fig. 3D).

**Fig. 3.**
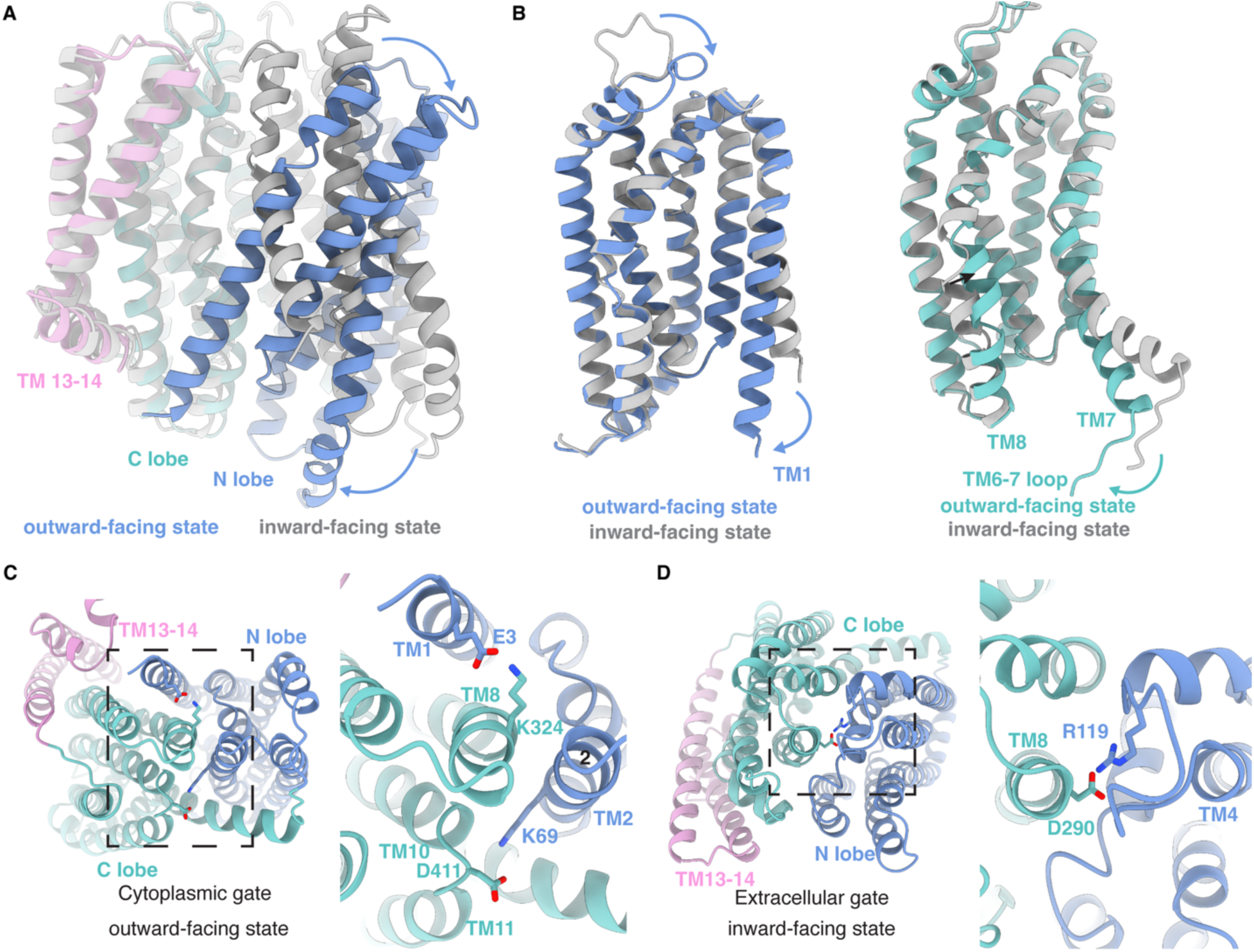
Conformational transitions during the lipid II transport cycle. (**A**) Structural overlay of inward-facing and outward-facing *Sa*MurJ aligned on the C-lobe. Arrows highlight the rigid-body movement of the N-lobe during inward-to-outward conformational transitions. Outward-facing *Sa*MurJ is colored as in Fig. 1 and inward-facing *Sa*MurJ is shown in gray. (**B**) Alignment of individual lobes colored as in (A). Arrows indicate the structural changes observed in TM1 of the N-lobe and TM7 of the C-lobe between the inward-facing and outward-facing conformations. (**C**) The cytoplasmic gate of the outward-facing state with the salt bridges at the gate highlighted. Interacting residue side chains are shown in stick representation and highlighted in the zoomed-in view on the right. (**D**) The extracellular gate of the inward-facing state, shown similar to (C).

### Predicted substrate-bound model in monoderm MurJ

To gain insight into the structural basis of lipid II recognition by *Sa*MurJ, we generated computational predictions of *Sa*MurJ bound to lipid II-Gly5, the *S. aureus*-specific form of lipid II (27, 28), using Chai-1 (29) (Fig. 4A). This predicted structure resembles our experimentally determined inward-facing conformation, which is a compatible state of MurJ to engage the lipid II substrate from the cytoplasm. Here, the undecaprenyl tail extends out from the membrane portal between TM1 and TM8 into the hydrophobic groove formed by TM13-14. The diphosphate moiety of the lipid II headgroup is positioned at the strongly cationic site adjacent to the membrane portal and groove (proximal site) (3) and coordinated by the conserved K22-R176-K302 triad (Fig. 4B). The predicted model places R153 and E172 near the disaccharide moiety of the lipid II headgroup, consistent with their previously reported functional importance and roles in substrate binding (15). The pentapeptide moiety is positioned at the distal site (far from the groove) in the C-lobe. The overall positioning of lipid II in the predicted structure is in general agreement with previous docking studies of lipid II to *Thermosipho africanus* MurJ (3). The additional penta-glycine modification of the pentapeptide moiety, specific to *S. aureus* lipid II, further extends away from the proximal site and contacts residues Q157 and E169 (Fig. 4B).

**Fig. 4.**
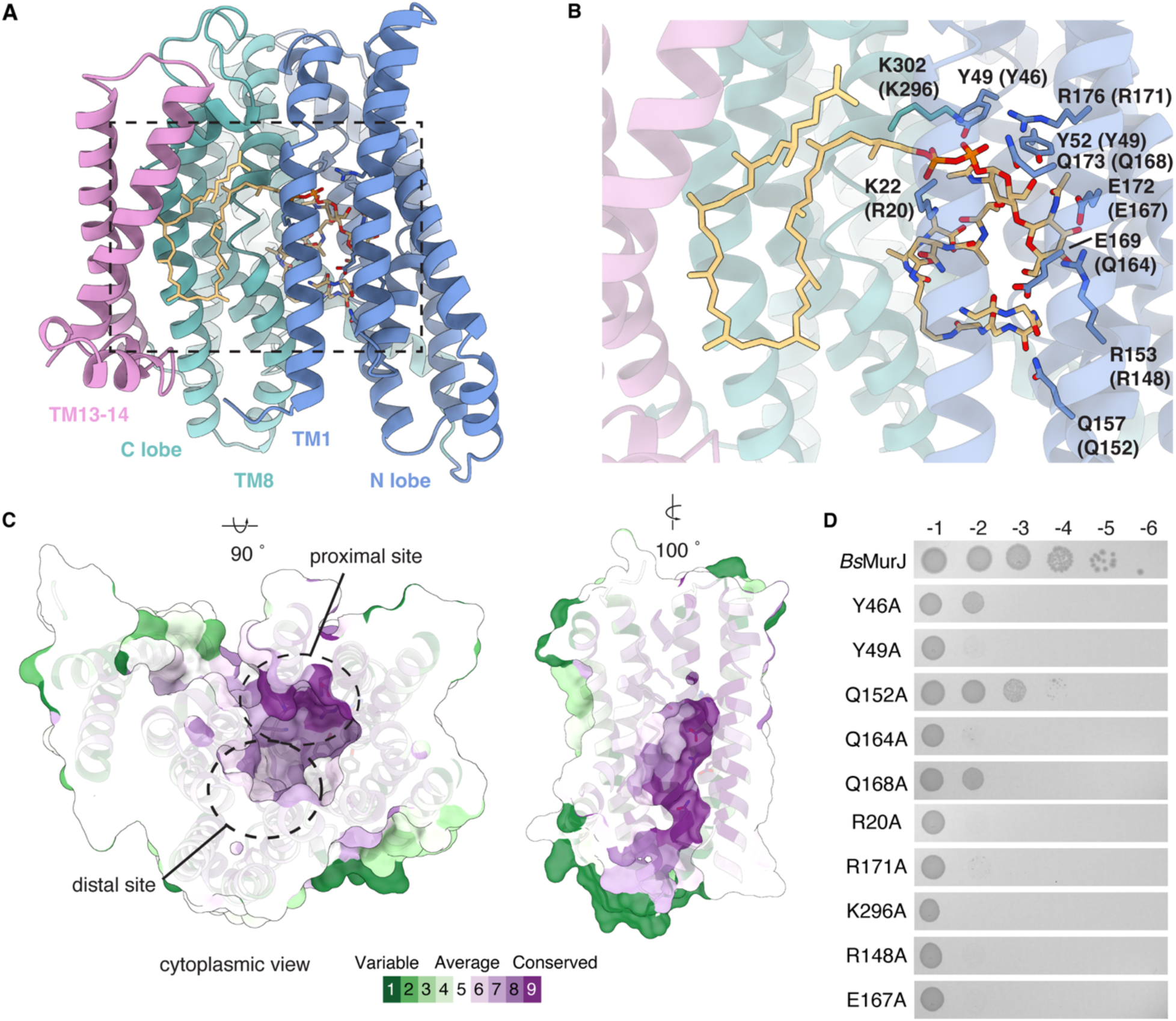
Model for substrate recognition in monoderm MurJ. (**A**) Chai-predicted co-structure of *Sa*MurJ bound to lipid II-Gly5 in an inward-facing state. *Sa*MurJ is colored by domain architecture as in Fig. 1. **(B)** A zoomed-in view of (A) highlighting the lipid II-binding pocket. Highly conserved residues in *Sa*MurJ within the substrate-binding pocket (Consurf grade 9 and 8) are shown as sticks and labeled, corresponding residues in *Bs*MurJ are indicated in parentheses. (**C**) Clipped surface representation of the predicted lipid II-binding pocket mapped onto the inward-facing structure of *Sa*MurJ colored according to conservation as calculated by Consurf (30). Left, proximal and distal sites viewed from the cytoplasm. Right, surface representation of the highly conserved diphosphate- and disaccharide-binding pocket in the N-lobe. (**D**) Complementation by *Bs*MurJ variants in a MurJ-depletion *E. coli* strain. The chromosomal *murJ_Ec_* is controlled by an arabinose-inducible promoter. *Bs*MurJ variants are expressed from an IPTG-inducible P*_lac_* promoter.

We mapped residue conservation scores from a subset of representative monoderm MurJ homologs onto our inward-facing structure (Fig. 4C). Residues forming the diphosphate- and disaccharide-binding pocket in the N-lobe, including those at the proximal site, are highly conserved. In contrast, the pentapeptide-binding distal site is comparatively less conserved. To probe the functional importance of these highly conserved substrate-binding residues, we sought to introduce alanine substitutions at these positions and test for complementation in *E. coli* strains depleted of endogenous MurJ (Fig. 4B). We first tested complementation by wild-type *Sa*MurJ and found that *Sa*MurJ failed to complement the MurJ-depleted *E. coli* strains (Fig. S8A). We analyzed the membrane fractions by western blotting and confirmed that *Sa*MurJ was expressed in the membrane (Fig. S7B). We therefore turned to the *Bacillus subtilis* MurJ homolog, which has previously been shown to complement MurJ depletion (12), as a representative monoderm model to test the functional importance of predicted substrate-binding residues. Alanine substitutions at these conserved residues within the predicted substrate-binding pocket abolished complementation, except for Q152A, which retained partial complementation (Fig. 4D). These results indicate that the conserved residues are important for MurJ function, which is consistent with their proposed role in binding the diphosphate and disaccharide moieties of lipid II. All mutants were detected in the membrane at levels comparable to the wild-type *Bs*MurJ sequence (Fig. S8, C and D), suggesting that the loss of complementation reflects impaired MurJ activity.

## Discussion

The lipid II flippase MurJ catalyzes the essential translocation of lipid II across the cytoplasmic membrane during peptidoglycan biosynthesis. MurJ is conserved across peptidoglycan-producing bacteria but exhibits substantial sequence divergence between diderm and monoderm bacteria. While structures of MurJ from diderm bacteria have been captured in multiple conformational states that support the alternating-access mechanism of lipid II transport, the structural basis of lipid II flipping in monoderm bacteria has remained unknown. Here we present structures of *Staphylococcus aureus* MurJ in both inward- and outward-facing conformations, providing the first structural view of a monoderm lipid II flippase. These structures show that monoderm MurJ retains the conserved MurJ fold and undergoes similar inward-to-outward conformational transitions. These findings place MurJ within the broader context of lipid-linked oligosaccharide flippases from the MOP superfamily, including bacterial Wzx-family transporters in O-antigen biosynthesis, the teichoic acid precursor flippase TacF, and the eukaryotic flippase Rft1 in N-linked glycosylation. Despite substantial divergence in sequence and substrate specificity, these transporters share a conserved structural fold, as evidenced by structures of TacF (31) and WzxE (32). The inward- and outward-facing states observed in *Sa*MurJ support a shared alternating-access transport mechanism, suggesting that conformational cycling is a general strategy employed by divergent flippases to translocate lipid-linked glycans across membranes.

Our structures reveal both conserved and divergent structural features between monoderm and diderm MurJ. Despite low sequence homology, key functional features are preserved. Notably, a strongly cationic site, formed by a triad of positively charged residues adjacent to the membrane portal, has been proposed to coordinate the diphosphate moiety of the lipid II headgroup. In *Sa*MurJ, the K22-R176-K302 triad occupies positions structurally equivalent to the R18-R24-R270 triad in *Ec*MurJ. Although R176 in *Sa*MurJ aligns with M171 in *Ec*MurJ, its side chain occupies the same region as R24 and serves an equivalent role in maintaining the local electrostatic environment of the putative diphosphate-binding site. The binding of lipid II to this cationic triad has been proposed to induce conformational transitions upon substrate recognition in diderm MurJ (17). Consistent with this model, we show that mutation of R176 traps MurJ in an inward-facing conformation, supporting that the cationic triad plays a key role in coupling substrate engagement to conformational transitions during the transport cycle and that this mechanism is conserved across diderm and monoderm MurJ.

Beyond this conserved putative diphosphate-binding pocket, *Sa*MurJ has several structural features that differ from those in *Ec*MurJ. In particular, *Sa*MurJ lacks the conserved G/A-E-G-A motif found in diderm MurJ homologs. In diderm MurJ, this motif introduces a helix break within TM2 that enables TM bending during conformational transitions of the transport cycle (3–5). The absence of the G/A-E-G-A motif in *Sa*MurJ suggests that monoderm MurJ likely relies on alternative structural features in TM2 to support these conformational transitions. This is likely achieved by the conserved residues G62 and P64 in TM2, which are positioned to function as a hinge that accommodates TM2 bending during conformational transitions. These findings suggest that while the overall transport mechanism is conserved, monoderm MurJ has evolved distinct structural features to achieve the equivalent functional transitions.

Additional differences are observed within the central cavity. *Sa*MurJ contains two charged residues in the central cavity (R153 and E172) that are important for function and are positioned near the disaccharide moiety of the lipid II headgroup in the predicted model, representing a distinct feature of monoderm MurJ that may contribute to substrate recognition. Using the computationally-predicted model, together with conservation analysis, we identified an additional series of conserved residues (Y49, Y52, Q157, E169 & Q173) lining the substrate-binding pocket in *Sa*MurJ. These residues are unique to monoderm MurJ homologs and highlight potential differences in substrate recognition.

The conserved and distinct structural features identified here represent tractable targets for inhibitors. Our comparative structural analysis provides a basis for both broad-spectrum and selective inhibition of MurJ. Specifically, conserved structural elements, such as the cationic diphosphate-binding site, can be exploited to enable broad-spectrum inhibition of MurJ across both diderm and monoderm bacteria. In contrast, structural features unique to monoderm or diderm MurJ may provide opportunities for selective inhibition. Diderm MurJ homologs, particularly those from ESKAPE pathogens and the World Health Organization (WHO) priority pathogens, are highly conserved (Fig. S9A). Monoderm MurJ homologs from priority pathogens are more broadly distributed and exhibit greater sequence diversity. This divergence may reflect differences in lipid II composition across monoderm bacteria, which vary substantially in the peptide stem and interpeptide bridge, in contrast to the comparatively conserved lipid II in diderm bacteria (1, 33). Our structures provide a roadmap for developing both broad-spectrum and selective antibiotics that target a critical component of cell wall biogenesis.

## Materials and Methods

### Expression and purification of *Sa*MurJ and *Ec*MurJ

pETDuet vector containing *S. aureus* MurJ^1-553^ (WT or R176A mutant) with a C-terminal ALFA-10xHis tag (*Sa*MurJ_ALFA_), *S. aureus* MurJ^8-553^ with an N-terminal BRIL fusion domain and a C-terminal 10xHis tag (*Sa*MurJ_BRIL_), or *E. coli* MurJ^1-500^ with a C-terminal ALFA-10xHis tag (*Ec*MurJ_ALFA_) was transformed into *E. coli* NiCo21 (DE3) cells (New England Biolabs). Colonies from the plate were used to inoculate a small starter culture in 50 mL Lysogeny Broth (LB) with 100 µg/mL Ampicillin. The starter culture was grown under shaking conditions at 225 rpm and 37 °C for 2 h, then used to inoculate 6 L of 2x Yeast Extract Tryptone medium with 100 µg/mL Ampicillin. Cells were grown at 37 °C with shaking until reaching an optical density (OD_600_) of 0.6-0.7 and were transferred to 18 °C for 30 min. Protein expression was induced by adding 400 μM Isopropyl β-D-1-thiogalactopyranoside (IPTG) and the culture was grown overnight (approximately 16 h). Cells were harvested by centrifugation at 9,000*g* for 10 min at 4 °C. Cell pellets were frozen and stored at -80 °C. All subsequent purification steps were performed at 4 °C. Thawed cell pellets were re-suspended in lysis buffer (20 mM Tris-HCl pH 8.0, 300 mM NaCl, 5% glycerol), homogenized, and lysed using a M-110L microfluidizer (Microfluidics). Unlysed cells and cell debris were removed by centrifugation at 23,500*g* for 30 min. The resulting supernatant was then centrifuged again at 168,000*g* for 1 h to isolate the membrane. Membrane pellets were flash frozen in liquid nitrogen and stored at -80 °C. The membrane pellet was resuspended and solubilized for 1 h in extraction buffer (20 mM HEPES pH 8.0, 300 mM NaCl, 5% glycerol, 1% (w/v) lauryl maltose neopentyl glycol (LMNG, Anatrace) and 10 mM imidazole). Unextracted membrane proteins were cleared by centrifugation at 168,000*g* for 30 min. The supernatant was incubated with 1 ml of Ni-NTA agarose resin (Qiagen) for 1 h and loaded onto a gravity column. The resin was washed with 50 column volumes of wash buffer (20 mM HEPES pH 8.0, 300 mM NaCl, 5% glycerol, 0.005% LMNG, 30 mM imidazole). The bound protein was eluted with 5 column volumes of elution buffer (20 mM HEPES pH 8.0, 150 mM NaCl, 5% glycerol, 0.005% LMNG, and 300 mM imidazole). Eluted protein was concentrated to approximately 500 µL using a 30 kDa MWCO Centrifugal filter (Amicon) and subjected to a Superdex 200 Increase 10/300 GL size exclusion chromatography column (Cytiva) pre-equilibrated with 20 mM HEPES pH 8.0, 150 mM NaCl, 5% glycerol, 0.005% LMNG. Fractions were analyzed by SDS-PAGE. For purifications of *Sa*MurJ_ALFA_ WT and *Sa*MurJ_BRIL_, cholesteryl hemisuccinate (CHS) was included as an additive detergent in all steps at a 10:1 LMNG/CHS ratio by mass.

### Preparation of *Sa*MurJ and *Ec*MurJ samples for cryo-EM

The anti-BRIL BAG2 Fab and anti-Fab Nb were purified as previously described (23, 34). The anti-ALFA Nb and the SENP^EuB^ protease were purified as previously described with minor modifications (35). The pTP298_His–Avi–SUMOEu_ALFA Nb plasmid (Addgene #199390) was expressed in Rosetta-gami2 competent cells (Novagen). Anti-ALFA Nb was affinity purified by Ni-NTA chromatography. The Ni-NTA resin was washed with 50 column volumes of 20 mM HEPES pH 8.0, 150 mM NaCl, 5% glycerol, and 30 mM imidazole. The resin was then incubated with 5 column volumes of 20 mM HEPES pH 8.0, 150 mM NaCl, 5% glycerol, and 1.2 µM SENP^EuB^ protease at 4 °C for 40 min to elute the anti-ALFA Nb. The SENP^EuB^ cleaved Nb was further purified through a Superdex 200 Increase 10/300 size exclusion chromatography column (Cytiva) in 20 mM HEPES pH 8.0, 150 mM NaCl, 5% glycerol.

Purified *Sa*MurJ_ALFA_ WT, R176A mutant, *Ec*MurJ_ALFA_ were incubated with 1.5-fold excess of anti-ALFA Nb at 4 °C for 1 h. Purified *Sa*MurJ_BRIL_ Sample was incubated with 1.2-fold and 1.4-fold molar excess of BAG2 Fab and elbow Nb at 4 °C for 1 h. Excess Fab or Nb were removed via size-exclusion chromatography using a Superdex S200 Increase 10/300 GL column (Cytiva). Peak fractions containing Fab or Nb bound MurJ were pooled and concentrated with a 100 kDa MWCO Centrifugal filter (Amicon) to ∼3-4 mg/mL for cryo-EM grids.

### Cryo-EM grid preparation and data collection

0.05% Fluorinated Octyl Maltoside (FOM, Anatrace) was added prior to vitrification. 2 µl of Sample was applied to a glow-discharged holey carbon grid (Quantifoil R1.2/1.3 Cu300 or Au 300). The grid was blotted at 4 °C, 100% humidity for 5.5 s at a blot force of +7 before plunging into liquid ethane cooled by liquid nitrogen using the FEI Vitrobot Mark IV.

Cryo-EM data were acquired at the Caltech cryo-EM Facility on a Thermo Fisher Scientific Titan Krios operated at an acceleration voltage of 300 keV and equipped with an energy filter and a K3 direct electron detector (Gatan) in correlated double Sampling mode. Datasets were collected at a nominal magnification of 130,000x, corresponding to pixel size of 0.325 Å. Movies were recorded in SerialEM in 40 frames with a total exposure dose of 70 e^-^/Å^2^ at a defocus range of −1.0 to −3.0 μm.

### Image processing, model building and refinement

All cryo-EM data processing was performed in cryoSPARC (36) (v4.6.1). Movies were gain-normalized and motion correction using Patch Motion Correction. Contrast transfer function (CTF) was estimated in Patch CTF in cryoSPARC. Micrographs were manually curated to remove low quality images based on CTF estimations, total frame motion and relative ice thickness. Initial round of particle picking was performed using blob picker tool (particle diameter 70-140 Å for MurJ_ALFA_ or 80-160 Å for MurJ_BRIL_). Particles were extracted with a 480-pixel box size for MurJ_ALFA_ or 512-pixel box size for MurJ_BRIL_ and classified through iterative rounds of 2D classification to remove featureless particles. Initial 3D models were generated using ab-initio reconstruction. Particles were classified and filtered through iterative of ab-initio reconstruction to remove particles in poorly resolved classes. 40,000-80,000 particles that resulted in cryo-EM maps with well resolved TMDs were selected for Topaz particle picking training (37). Particle picked by Topaz were subject to one round of 2D classification. Particles were classified by multiple rounds of 3D classification via 2-class ab-initio reconstruction. Final sets of particles were polished through reference-based motion correction after an initial round of non-uniform refinement. Polished particles were refined by another round of non-uniform refinement (38) and further refined by local refinement with a soft mask covering the TM region.

For all model building and refinement, maps from focused local refinement were used. A predicted model of *Sa*MurJ was generated using AlphaFold (39), which produced an inward-facing state. An initial model of *Sa*MurJ in an outward-facing state was built by docking the N and C lobes of the AlphaFold model of *Sa*MurJ separately into the cryo-EM map as rigid bodies in UCSF ChimeraX. *Ec*MurJ from the previously determined Sgl^M^-MurJ complex structure (PDB:9NU4) was used as an initial model to build into the cryo-EM map. Manual model building was then performed in Coot (40). The models were refined through iterative rounds manual adjustments in Coot and real-space refinement in Phenix (41). The final structural models were visually inspected and validated in Molprobity (42).

### Evolutionary conservation analysis

Homologous sequences of MurJ were collected using JackHMMER (43) with the full-length *Staphylococcus aureus* MurJ sequence as the query. Searches were performed using a significance E-value threshold of 0.001. Sequences with lengths outside ±15% of the query were excluded. The remaining sequences were clustered using CD-HIT (44) with a sequence identity cutoff of 45% and a minimum alignment coverage of 80% to reduce redundancy while preserving sequence diversity. The resulting dataset of 244 homologs was aligned using MAFFT (45). Evolutionary conservation scores were calculated using Consurf (30) using the default settings. The Chai-1 predicted substrate-bound *Sa*MurJ model was used as the structural template. Residue conservation grades (1–9) were mapped onto the model and visualized in ChimeraX (46).

### Complementation by MurJ homologs

*Sa*MurJ and *Bs*MurJ was cloned into plasmid pCS126 (16) (*cat lacI* P*_lac_::murJ^+^-flag*) replacing the wild-type *E. coli murJ*. Plasmids encoding *Sa*MurJ and *Bs*MurJ variants were transformed into a conditional MurJ-depletion strain CS7 (2). Endogenous MurJ expression in this strain is under the control of an arabinose-inducible promoter (P*_BAD_*). Transformants of CS7 were selected on Lysogeny Broth (LB) agar supplemented with 25 µg/mL ampicillin, 34 µg/mL chloramphenicol, and 0.2% arabinose. Cells were transferred to 5 mL of LB medium with 25 µg/mL ampicillin, 34 µg/mL chloramphenicol and 0.2% arabinose. Cultures were diluted and normalized to an optical density (OD_600_) of 0.8. Serial dilutions (10^-1^ to 10^-5^) of the normalized cell cultures were spotted on LB agar plates supplemented with either 0.2% arabinose or 100 µM Isopropyl β-D-1-thiogalactopyranoside (IPTG). Plates were imaged after incubation at 37 °C for 16 h.

### Detection of MurJ variants in the membrane by Western blotting

Plasmids encoding MurJ variants were transformed into CS7 (2). Colonies from the plate were inoculated into a 5 mL LB culture with 325 µg/mL ampicillin, 34 µg/mL chloramphenicol, and 0.2% arabinose. The cultures were grown shaking at 225 rpm and 37 °C. When OD_600_ reached ∼0.8, expression of the plasmid-borne MurJ variants was induced with the addition of 0.4 mM IPTG. Cells were harvested at 1 h post-induction by centrifugation at 4,000*g* for 10 minutes at 4 °C. Cell pellets were resuspended in 20 mM Tris-HCl pH 8.0, 300 mM NaCl, and 5% glycerol in volumes normalized to final OD_600_ (50 uL per OD_600_ units). Resuspended cell pellets were kept on ice and lysed using the Misonix S-4000 Sonicator with microtip (amplitude of 20, 1 min cycle, 10 s ON, 2 s OFF). Unlysed cells and cell debris were removed by centrifugation at 213,000*g* for 20 min at 4 °C. The supernatant was then centrifuged to collect the membrane fraction using the Optima TLX Tabletop Ultracentrifuge (Beckman Coulter) at 108,628*g* for 30 min at 4 °C in TLA100 rotor. Membrane pellets were resuspended in 30 uL of 20 mM HEPES pH 8.0, 150 mM NaCl and 0.01% LMNG. 10 µL of the membrane fraction samples were resolved on pre-cast Any kD gels (Bio-Rad) and transferred to a nitrocellulose membrane using standard semi-dry transfer protocols. Total protein on the membrane was visualized on a using Revert 700 Total Protein Stain (LiCor). Membranes were blocked with 5% milk in Tris-buffered saline with Tween 20 for 1 h at room temperature and then probed with anti-His primary antibodies (Sigma-Aldrich) at 1:2,000 dilution at 4 °C overnight and anti-Mouse-IRDye 800CW secondary antibodies (LiCor) at 10,000 dilutions for 1 h at room temperature. Blots were imaged using the Odyssey LiCor infrared Imager. Protein levels in the membrane were normalized to total protein levels and quantified using Empiria Studio (LiCor).

## Acknowledgments

(Cryo)Electron microscopy was performed in the Beckman Institute Resource Center for Transmission Electron Microscopy at Caltech. We thank Songye Chen and Tyler Brittain for assistance with cryo-EM data collection. We thank the Voorhees lab at Caltech for the kind gift of the pTP298 ALFA Nb plasmid. We thank Doug Rees for comments on the manuscript.

Funding for this work was provided by National Institutes of Health grant R01GM114611 (W.M.C.), the G. Harold and Leila Y. Mathers Foundation (W.M.C.), and the Chan Zuckerberg Initiative.

## Data and materials availability

Cryo-EM maps have been deposited into in the Electron Microscopy Data Bank (EMDB) under accession codes EMD-76175 (*Sa*MurJ in an outward state), EMD-76177 (*Sa*MurJ in an inward state), EMD-76178 (*Sa*MurJ R176A mutant), and EMD-76176 (*Ec*MurJ in an outward state). Coordinates for the atomic models have been deposited in the Protein Data Bank (PDB) under accession codes 11XX (*Sa*MurJ in an outward state), 11XZ (*Sa*MurJ in an inward state), 11YA (*Sa*MurJ R176A mutant), and 11XY (*Ec*MurJ in an outward state).

## References

1. S. Kumar, A. Mollo, D. Kahne, N. Ruiz, The Bacterial Cell Wall: From Lipid II Flipping to Polymerization. Chemical Reviews 122, 8884–8910 (2022).

2. L.-T. Sham, et al., MurJ is the flippase of lipid-linked precursors for peptidoglycan biogenesis. Science 345, 220–222 (2014).

3. A. C. Y. Kuk, E. H. Mashalidis, S.-Y. Lee, Crystal structure of the MOP flippase MurJ in an inward-facing conformation. Nature Structural & Molecular Biology 24, 171–176 (2017).

4. S. Zheng, et al., Structure and mutagenic analysis of the lipid II flippase MurJ from Escherichia coli. Proceedings of the National Academy of Sciences 115, 201802192 (2018).

5. A. C. Y. Kuk, A. Hao, Z. Guan, S.-Y. Lee, Visualizing conformation transitions of the Lipid II flippase MurJ. Nature Communications 10, 1736 (2019).

6. H. Kohga, et al., Crystal structure of the lipid flippase MurJ in a “squeezed” form distinct from its inward- and outward-facing forms. Structure (2022). 10.1016/j.str.2022.05.008.

7. N. Ruiz, *Streptococcus pyogenes* YtgP (Spy_0390) Complements *Escherichia coli* Strains Depleted of the Putative Peptidoglycan Flippase MurJ. Antimicrob Agents Chemother 53, 3604–3605 (2009).

8. C. A. Caffalette, J. Kuklewicz, N. Spellmon, J. Zimmer, Biosynthesis and Export of Bacterial Glycolipids. Annual Review of Biochemistry 89, 741–768 (2020).

9. A. C. Y. Kuk, A. Hao, S.-Y. Lee, Structure and Mechanism of the Lipid Flippase MurJ. Annu Rev Biochem 91, 705–729 (2022).

10. T. Pomorski, A. K. Menon, Lipid flippases and their biological functions. Cell. Mol. Life Sci. 63, 2908–2921 (2006).

11. A. Fay, J. Dworkin, *Bacillus subtilis* Homologs of MviN (MurJ), the Putative *Escherichia coli* Lipid II Flippase, Are Not Essential for Growth. J Bacteriol 191, 6020–6028 (2009).

12. A. J. Meeske, et al., MurJ and a novel lipid II flippase are required for cell wall biogenesis in Bacillus subtilis. Proceedings of the National Academy of Sciences 112, 6437–6442 (2015).

13. A. S. Ramírez, K. P. Locher, Structural and mechanistic studies of the *N* -glycosylation machinery: from lipid-linked oligosaccharide biosynthesis to glycan transfer. Glycobiology 33, 861–872 (2023).

14. E. K. Butler, R. M. Davis, V. Bari, P. A. Nicholson, N. Ruiz, Structure-Function Analysis of MurJ Reveals a Solvent-Exposed Cavity Containing Residues Essential for Peptidoglycan Biogenesis in Escherichia coli. Journal of Bacteriology 195, 4639–4649 (2013).

15. E. K. Butler, W. B. Tan, H. Joseph, N. Ruiz, Charge Requirements of Lipid II Flippase Activity in Escherichia coli. Journal of Bacteriology 196, 4111–4119 (2014).

16. L. Sham, S. Zheng, A. A. Yakhnina, A. C. Kruse, T. G. Bernhardt, Loss of specificity variants of WzxC suggest that substrate recognition is coupled with transporter opening in MOP-family flippases. Molecular Microbiology 109, 633–641 (2018).

17. F. A. Rubino, et al., Detection of Transport Intermediates in the Peptidoglycan Flippase MurJ Identifies Residues Essential for Conformational Cycling. Journal of the American Chemical Society 142, 5482–5486 (2020).

18. J. E. Mott, et al., Resistance mapping and mode of action of a novel class of antibacterial anthranilic acids: evidence for disruption of cell wall biosynthesis. Journal of Antimicrobial Chemotherapy 62, 720–729 (2008).

19. J. Huber, et al., Chemical Genetic Identification of Peptidoglycan Inhibitors Potentiating Carbapenem Activity against Methicillin-Resistant Staphylococcus aureus. Chemistry & Biology 16, 837–848 (2009).

20. J. Chu, et al., Discovery of MRSA active antibiotics using primary sequence from the human microbiome. Nature chemical biology 12, 1004–1006 (2016).

21. J. Chu, et al., Human Microbiome Inspired Antibiotics with Improved β-Lactam Synergy against MDR Staphylococcus aureus. ACS Infectious Diseases 4, 33–38 (2018).

22. Y. E. Li, et al., Convergent MurJ flippase inhibition by phage lysis proteins. Nature (2026). 10.1038/s41586-026-10163-w.

23. S. Mukherjee, et al., Synthetic antibodies against BRIL as universal fiducial marks for single−particle cryoEM structure determination of membrane proteins. Nature Communications 11, 1598 (2020).

24. H. Götzke, et al., The ALFA-tag is a highly versatile tool for nanobody-based bioscience applications. Nature Communications 10, 4403 (2019).

25. S. Kumar, F. A. Rubino, A. G. Mendoza, N. Ruiz, The bacterial lipid II flippase MurJ functions by an alternating-access mechanism. Journal of Biological Chemistry 294, 981–990 (2019).

26. D. Drew, O. Boudker, Ion and lipid orchestration of secondary active transport. Nature 626, 963–974 (2024).

27. M. Matsuhashi, C. P. Dietrich, J. L. Strominger, Incorporation of glycine into the cell wall glycopeptide in Staphylococcus aureus: role of sRNA and lipid intermediates. Proc. Natl. Acad. Sci. U.S.A. 54, 587–594 (1965).

28. T. Schneider, et al., *In vitro* assembly of a complete, pentaglycine interpeptide bridge containing cell wall precursor (lipid II-Gly_5_) of *Staphylococcus aureus*. Molecular Microbiology 53, 675–685 (2004).

29. Chai Discovery, et al., Chai-1: Decoding the molecular interactions of life. [Preprint] (2024). Available at: http://biorxiv.org/lookup/doi/10.1101/2024.10.10.615955 [Accessed 25 March 2025].

30. H. Ashkenazy, et al., ConSurf 2016: an improved methodology to estimate and visualize evolutionary conservation in macromolecules. Nucleic Acids Res 44, W344–W350 (2016).

31. G. Cebrero, et al., Mechanistic basis of teichoic acid transport by a gatekeeper flippase. Nat Commun (2026). 10.1038/s41467-026-73616-w.

32. A. Le Bas, et al., Structure of WzxE the lipid III flippase for Enterobacterial Common Antigen polysaccharide. Open Biol. 15, 240310 (2025).

33. D. Münch, H.-G. Sahl, Structural variations of the cell wall precursor lipid II in Gram-positive bacteria — Impact on binding and efficacy of antimicrobial peptides. Biochimica et Biophysica Acta (BBA) - Biomembranes 1848, 3062–3071 (2015).

34. J. Ereño-Orbea, et al., Structural Basis of Enhanced Crystallizability Induced by a Molecular Chaperone for Antibody Antigen-Binding Fragments. Journal of Molecular Biology 430, 322–336 (2018).

35. T. A. Stevens, et al., A nanobody-based strategy for rapid and scalable purification of native human protein complexes. (2023). 10.1101/2023.03.09.531980.

36. A. Punjani, J. L. Rubinstein, D. J. Fleet, M. A. Brubaker, cryoSPARC: algorithms for rapid unsupervised cryo-EM structure determination. Nature Methods 14, 290–296 (2017).

37. T. Bepler, et al., Positive-unlabeled convolutional neural networks for particle picking in cryo-electron micrographs. Nat Methods 16, 1153–1160 (2019).

38. A. Punjani, H. Zhang, D. J. Fleet, Non-uniform refinement: adaptive regularization improves single-particle cryo-EM reconstruction. Nat Methods 17, 1214–1221 (2020).

39. J. Jumper, et al., Highly accurate protein structure prediction with AlphaFold. Nature 596, 583–589 (2021).

40. P. Emsley, B. Lohkamp, W. G. Scott, K. Cowtan, Features and development of *Coot*. Acta Crystallogr D Biol Crystallogr 66, 486–501 (2010).

41. P. V. Afonine, et al., Real-space refinement in *PHENIX* for cryo-EM and crystallography. Acta Crystallogr D Struct Biol 74, 531–544 (2018).

42. C. J. Williams, et al., MolProbity: More and better reference data for improved all-atom structure validation. Protein Science 27, 293–315 (2018).

43. S. C. Potter, et al., HMMER web server: 2018 update. Nucleic Acids Research 46, W200–W204 (2018).

44. W. Li, A. Godzik, Cd-hit: a fast program for clustering and comparing large sets of protein or nucleotide sequences. Bioinformatics 22, 1658–1659 (2006).

45. K. Katoh, J. Rozewicki, K. D. Yamada, MAFFT online service: multiple sequence alignment, interactive sequence choice and visualization. Briefings in Bioinformatics 20, 1160–1166 (2019).

46. T. D. Goddard, et al., UCSF ChimeraX: Meeting modern challenges in visualization and analysis. Protein Science 27, 14–25 (2018).

