## Supplemental Information for "The structure of the lipid II flippase from monoderm bacteria"

**This PDF file includes:**

Figs. S1 to S9
Tables S1
SI References

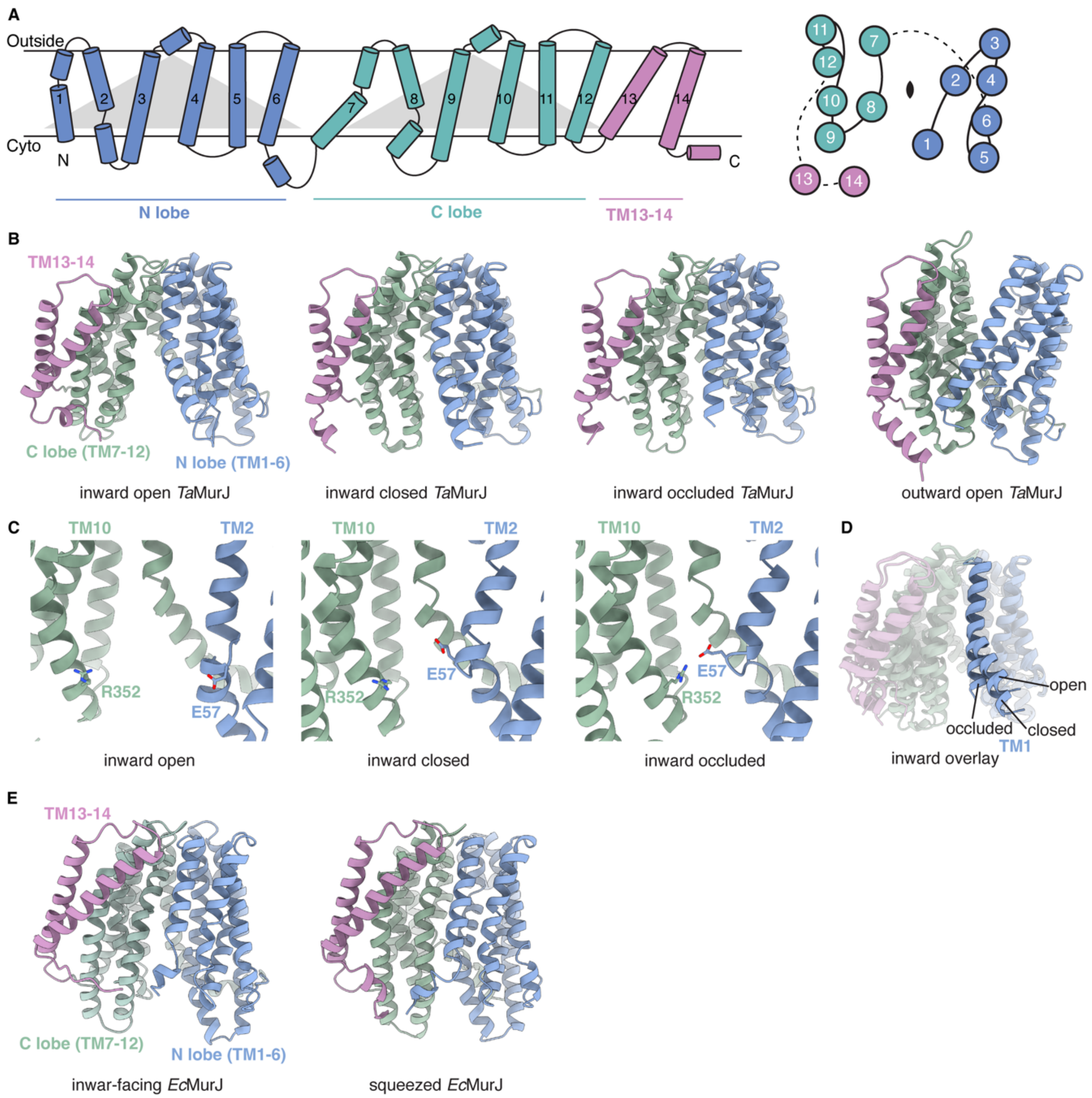


**Fig. S1. Architecture of MurJ. (A)** Left, predicted topology of *Sa*MurJ showing the 14 numbered transmembrane helices organized into the N lobe (TM1-6) (blue), C lobe (TM7-12) (green), and TM13-14 (pink). Triangles indicate the symmetrical lobes. Right, top view representation of MurJ showing the arrangement of transmembrane helices surrounding the central cavity. TMs1-12 are pseudo-C2-symmetric six-helix N- and C-lobes. **(B)** Crystal structures of *Thermosipho africanus* MurJ (*Ta*MurJ) **(1)** in distinct conformational states colored by domain architecture (PDB: 6NC7, 6NC6, 6NC8, 6NC9). **(C)** Close-up view of the thin gate between Glu57 in TM2 and R352 in TM10. **(D)** Structural overlay of inward structures of *Ta*MurJ highlighting the closure of TM1 at the membrane portal. **(E)** Crystal structures of *E. coli* MurJ (*Ec*MurJ) in an inward-facing state **(2)** (PDB: 6CC4) or a squeezed state **(3)** (PDB: 7WAG).

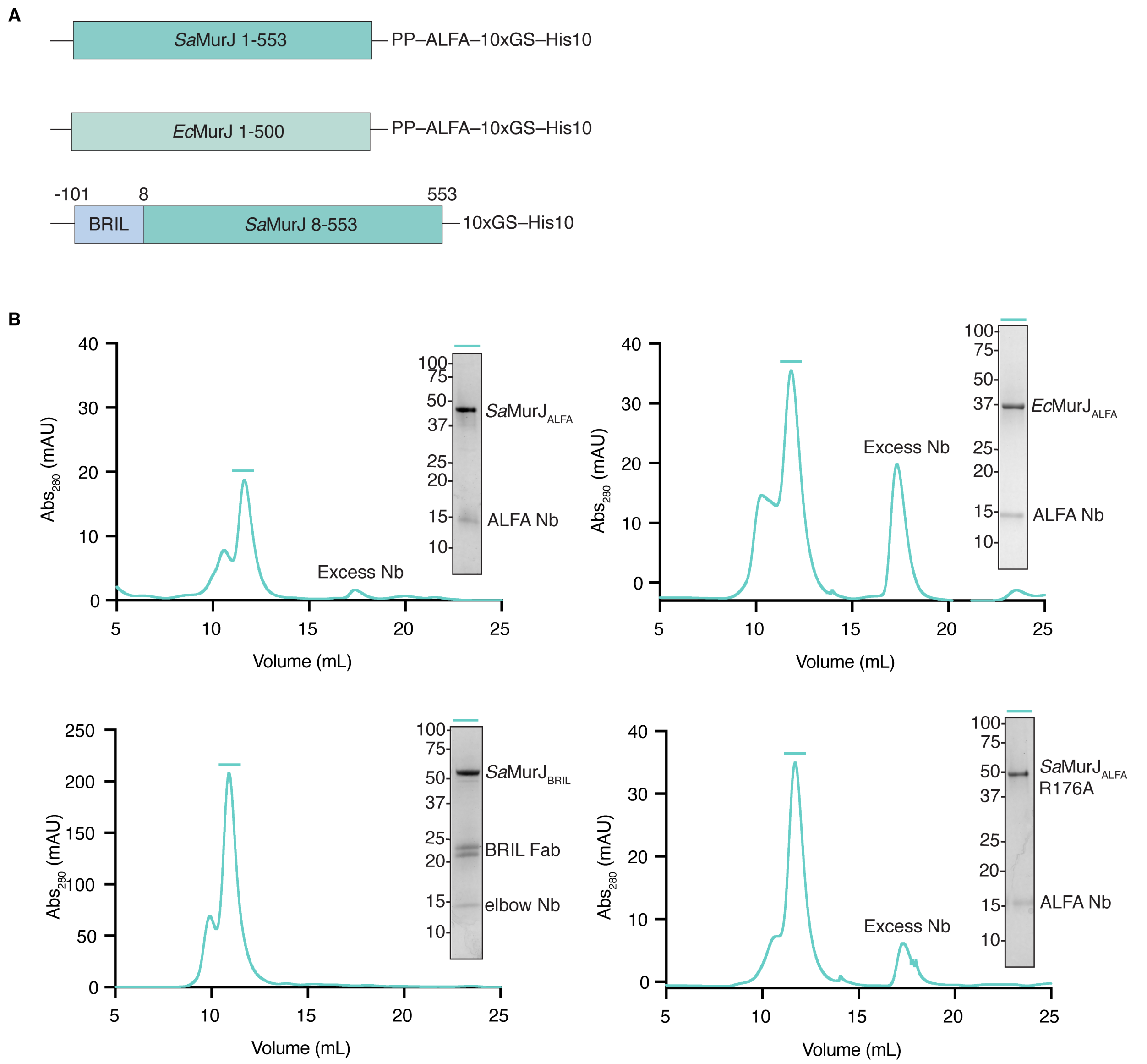


**Fig. S2. Construct design and purification of *Sa*MurJ and *Ec*MurJ proteins.** (**A**) Schematic of expression constructs for *Sa*MurJ and *Ec*MurJ. (**B**) Representative size-exclusion chromatography profiles and SDS-PAGE analyses of purified *Sa*MuJ$_{\text{ALFA}}$-ALFANb, *Sa*MurJ$_{\text{BRIL}}$-Fab-Nb, *Ec*MurJ$_{\text{ALFA}}$-ALFANb complexes.

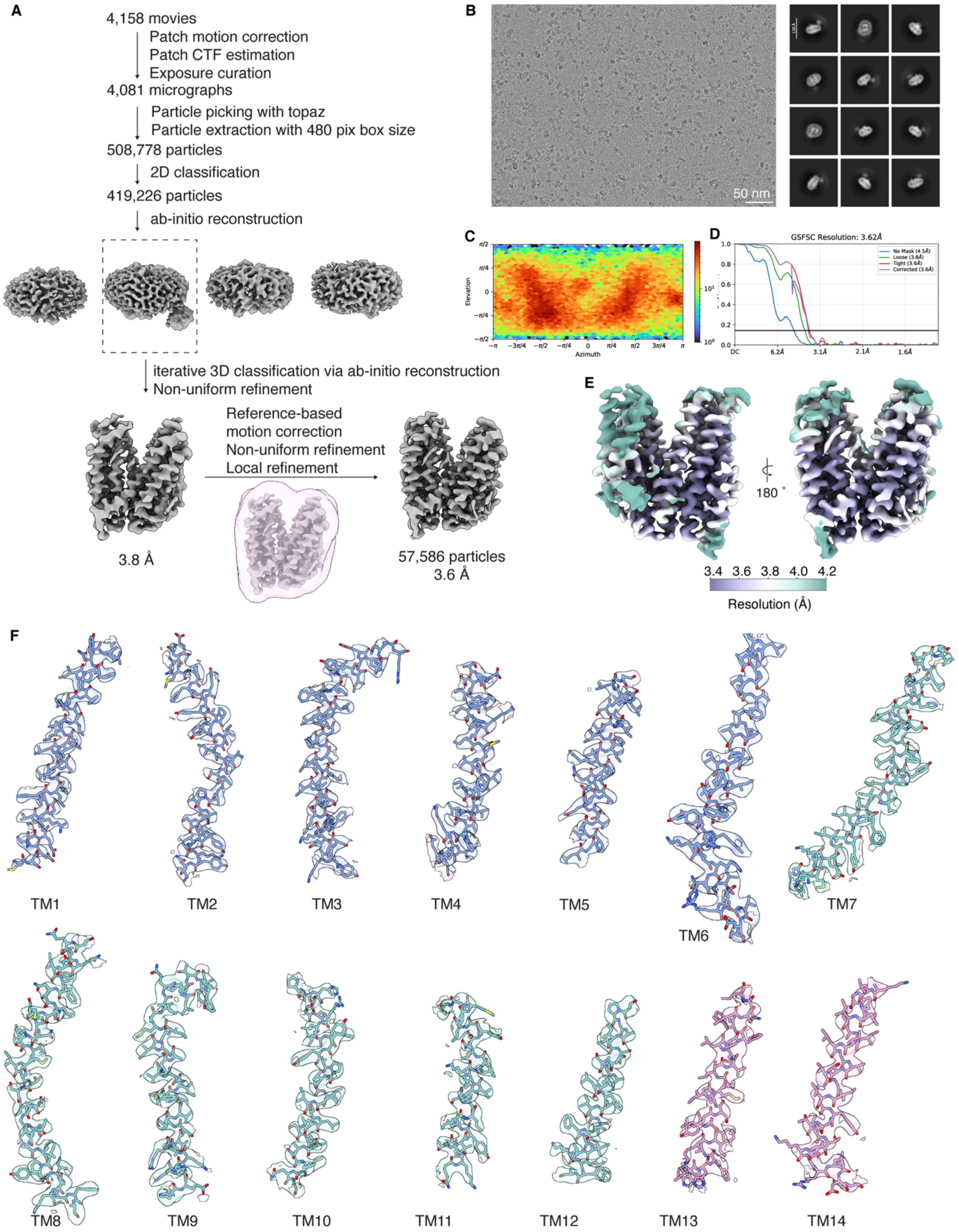
A
4,158 movies
Patch motion correction
Patch CTF estimation
Exposure curation
4,081 micrographs
Particle picking with topaz
Particle extraction with 480 pix box size
508,778 particles
2D classification
419,226 particles
ab-initio reconstruction
iterative 3D classification via ab-initio reconstruction
Non-uniform refinement
Reference-based
motion correction
Non-uniform refinement
Local refinement
3.8 Å
57,586 particles
3.6 Å
B
50 nm
C
Elevation
Azimuth
D
GSFSC Resolution: 3.62Å
No Mask (4.5Å)
Loose (3.8Å)
Tight (3.6Å)
Corrected (3.6Å)
E
180 °
3.4 3.6 3.8 4.0 4.2
Resolution (Å)
F
TM1
TM2
TM3
TM4
TM5
TM6
TM7
TM8
TM9
TM10
TM11
TM12
TM13
TM14

**Fig. S3. Cryo-EM structure determination of *Sa*MurJ in an outward-facing state.** (**A**) Summary of cryo-EM data processing workflow to obtain the outward-facing structure of *Sa*MurJ. (**B**) Representative micrograph (left) and 2D class averages (right). (**C**) Angular distribution of particles of the final 3D reconstruction. (**D**) Fourier shell correlation curve between two half maps. (**E**) Local resolution map of the final cryo-EM map colored by resolution from 3.4 Å (purple) to 4.2 Å (green). (**F**) Cryo-EM density of each TM helix of *Sa*MurJ. Map contour level = 0.059 in ChimeraX **(4)**.

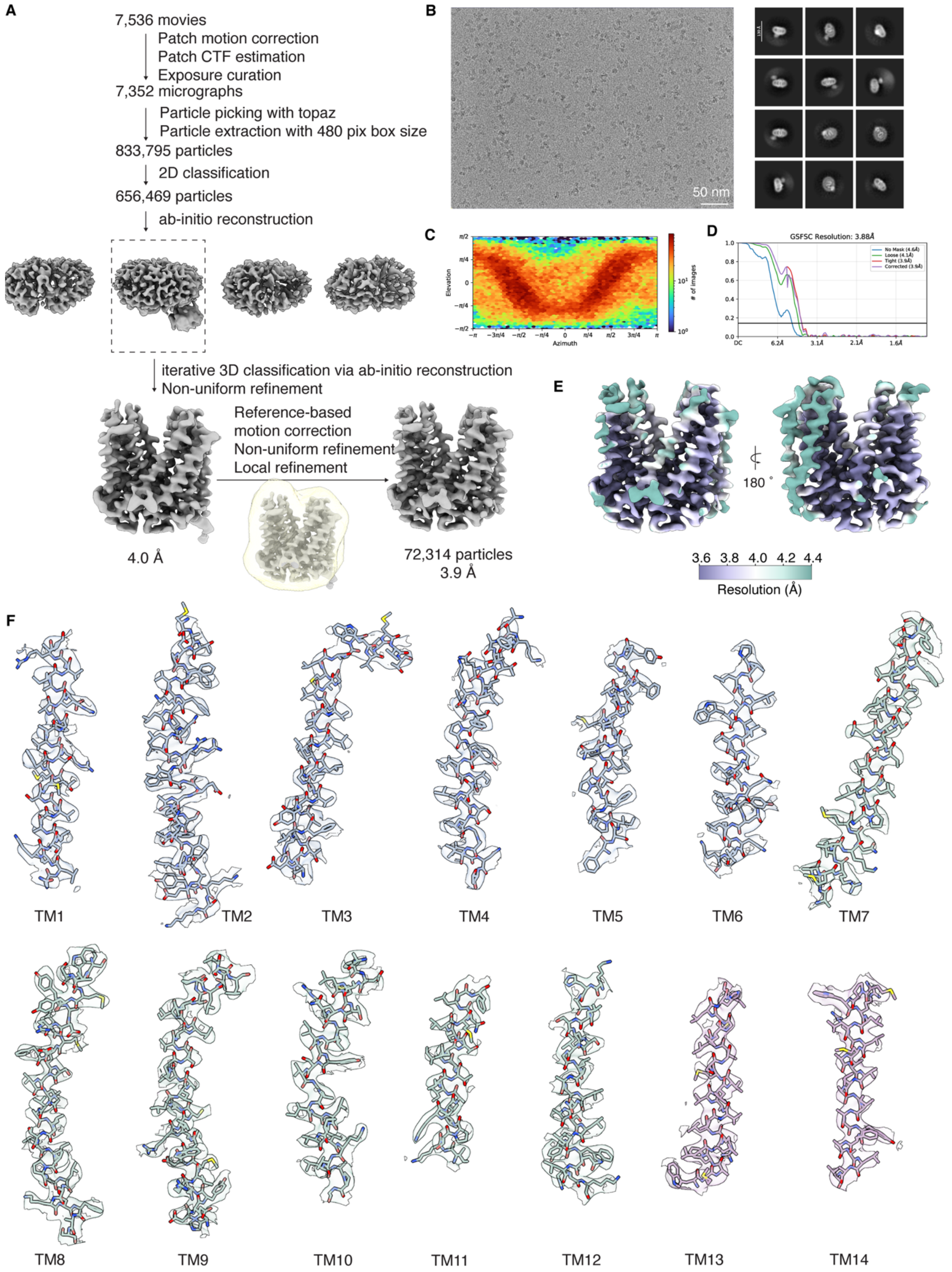
A
7,536 movies
Patch motion correction
Patch CTF estimation
Exposure curation
7,352 micrographs
Particle picking with topaz
Particle extraction with 480 pix box size
833,795 particles
2D classification
656,469 particles
ab-initio reconstruction
iterative 3D classification via ab-initio reconstruction
Non-uniform refinement
Reference-based
motion correction
Non-uniform refinement
Local refinement
4.0 Å
72,314 particles
3.9 Å
B
50 nm
C
Elevation
Azimuth
### of images
D
GSFSC Resolution: 3.88Å
No Mask (4.6Å)
Loose (4.1Å)
Tight (3.9Å)
Corrected (3.9Å)
E
180 °
3.6 3.8 4.0 4.2 4.4
Resolution (Å)
F
TM1
TM2
TM3
TM4
TM5
TM6
TM7
TM8
TM9
TM10
TM11
TM12
TM13
TM14

**Fig. S4. Cryo-EM structure determination of *Ec*MurJ in an outward-facing state.** (**A**) Summary of cryo-EM data processing workflow to obtain the outward-facing structure of *Ec*MurJ. (**B**) Representative micrograph (left) and 2D class averages (right). (**C**) Angular distribution of particles of the final 3D reconstruction. (**D**) Fourier shell correlation curves between two half maps. (**E**) Local resolution map of the final cryo-EM map colored by resolution from 3.6 Å (purple) to 4.4 Å (green). (**F**) Cryo-EM density of each TM helix of *Ec*MurJ. Map contour level = 0.055 in ChimeraX **(4)**.

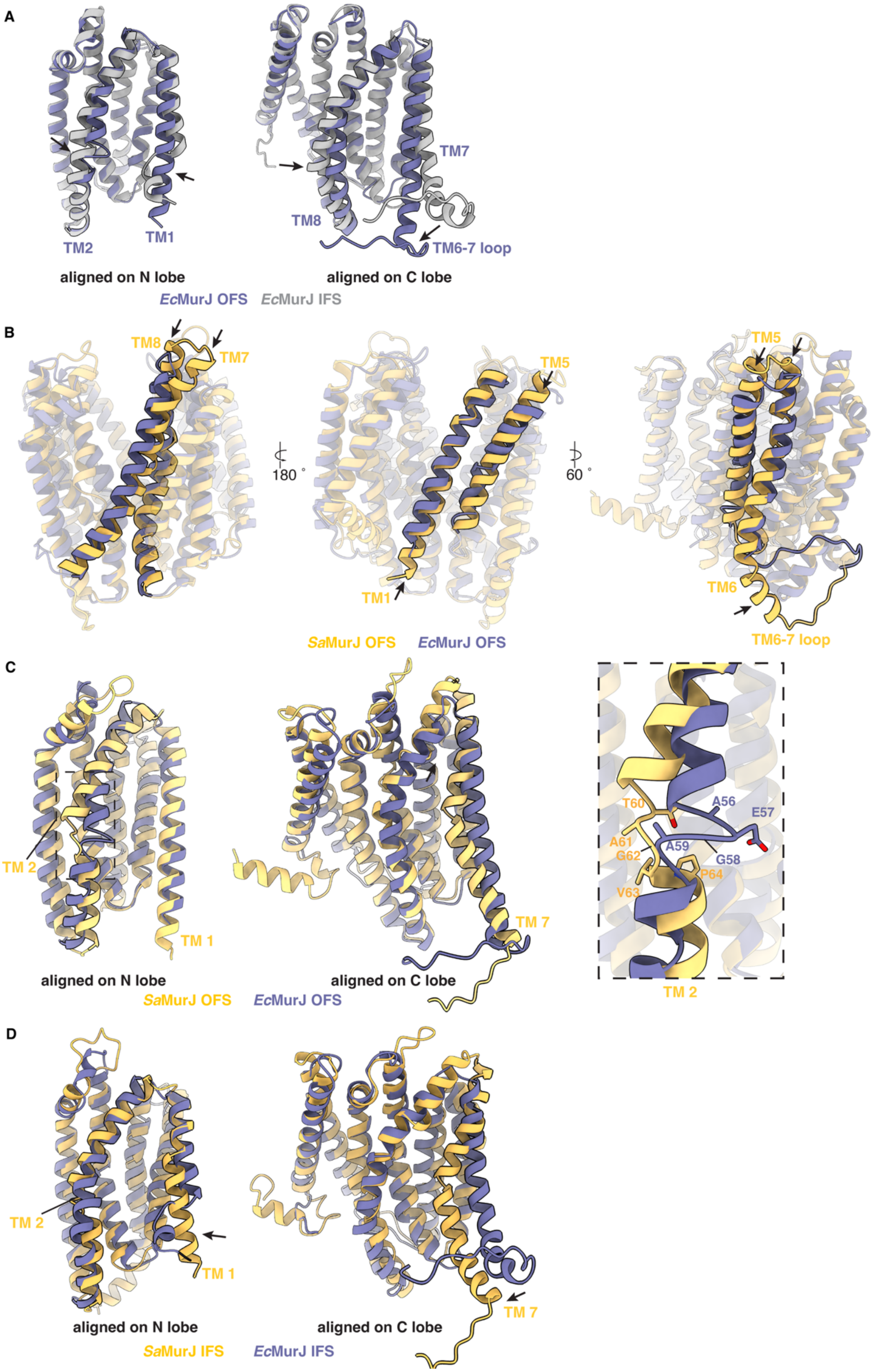
A
TM2
TM1
TM8
TM7
TM6-7 loop
aligned on N lobe
aligned on C lobe
EcMurJ OFS
EcMurJ IFS
B
TM8
TM7
TM5
TM1
TM6
TM6-7 loop
180 °
60 °
SaMurJ OFS
EcMurJ OFS
C
TM 2
TM 1
TM 7
T60
A61
G62
V63
P64
A56
E57
A59
G58
aligned on N lobe
aligned on C lobe
SaMurJ OFS
EcMurJ OFS
D
TM 2
TM 1
TM 7
aligned on N lobe
aligned on C lobe
SaMurJ IFS
EcMurJ IFS

**Fig. S5. Structural comparison of *Sa*MurJ and *Ec*MurJ. (A)** Structural overlay of the outward- and inward-facing *Ec*MurJ (PDB: 6CC4) aligned to either the N- or the C-lobe. (**B**) Structural overlay of outward-facing *Sa*MurJ and *Ec*MurJ highlighting the differences in TMs1, 5, 7, 8, and TM6-7 loop. (**C**) Comparison of the outward-facing *Sa*MurJ and *Ec*MurJ aligned to either the N- or the C-lobe. Dashed box highlights structural and sequence differences in TM2. (**D**) Comparison of the inward-facing *Sa*MurJ and *Ec*MurJ (PDB: 6CC4) aligned to either the N- or the C-lobe.

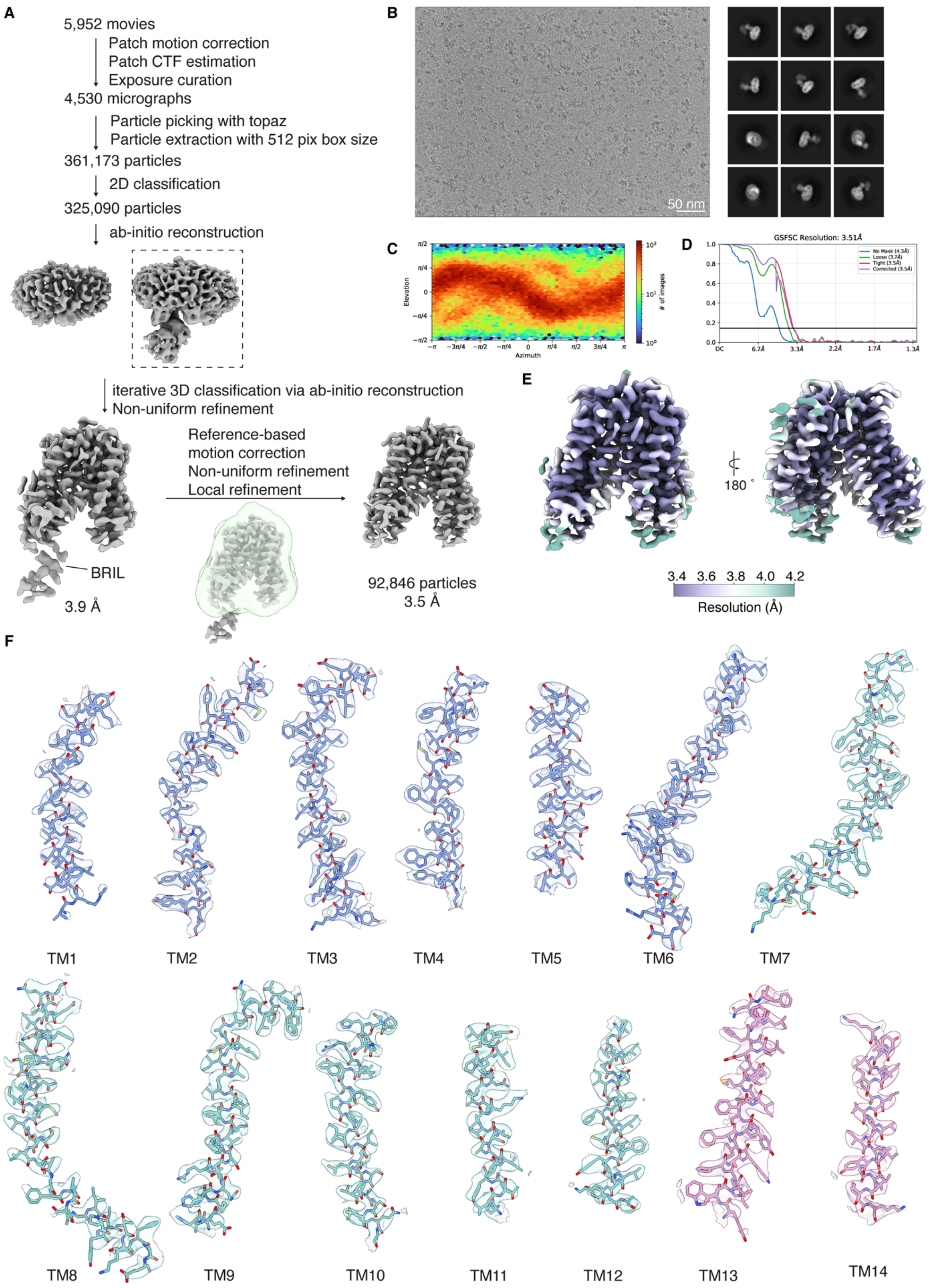

A
5,952 movies
Patch motion correction
Patch CTF estimation
Exposure curation
4,530 micrographs
Particle picking with topaz
Particle extraction with 512 pix box size
361,173 particles
2D classification
325,090 particles
ab-initio reconstruction
iterative 3D classification via ab-initio reconstruction
Non-uniform refinement
BRIL
3.9 Å
Reference-based
motion correction
Non-uniform refinement
Local refinement
92,846 particles
3.5 Å
B
50 nm
C
Elevation
Azimuth
### of images
D
GSFSC Resolution: 3.51Å
No Mask (4.3Å)
Loose (3.7Å)
Tight (3.5Å)
Corrected (3.5Å)
E
180 °
3.4 3.6 3.8 4.0 4.2
Resolution (Å)
F
TM1
TM2
TM3
TM4
TM5
TM6
TM7
TM8
TM9
TM10
TM11
TM12
TM13
TM14

**Fig. S6. Cryo-EM structure determination of *Sa*MurJ in an inward-facing state.** (**A**) Summary of cryo-EM data processing workflow to obtain the inward-facing structure of *Sa*MurJ. (**B**) Representative micrograph (left) and 2D class averages (right). (**C**) Angular distribution of particles of the final 3D reconstruction. (**D**) Fourier shell correlation curves between two half maps. (**E**) Local resolution map of the final cryo-EM map colored by resolution from 3.4 Å (purple) to 4.2 Å (green). (**F**) Cryo-EM density of each TM helix of *Sa*MurJ. Map contour level = 0.061 in ChimeraX **(4)**.

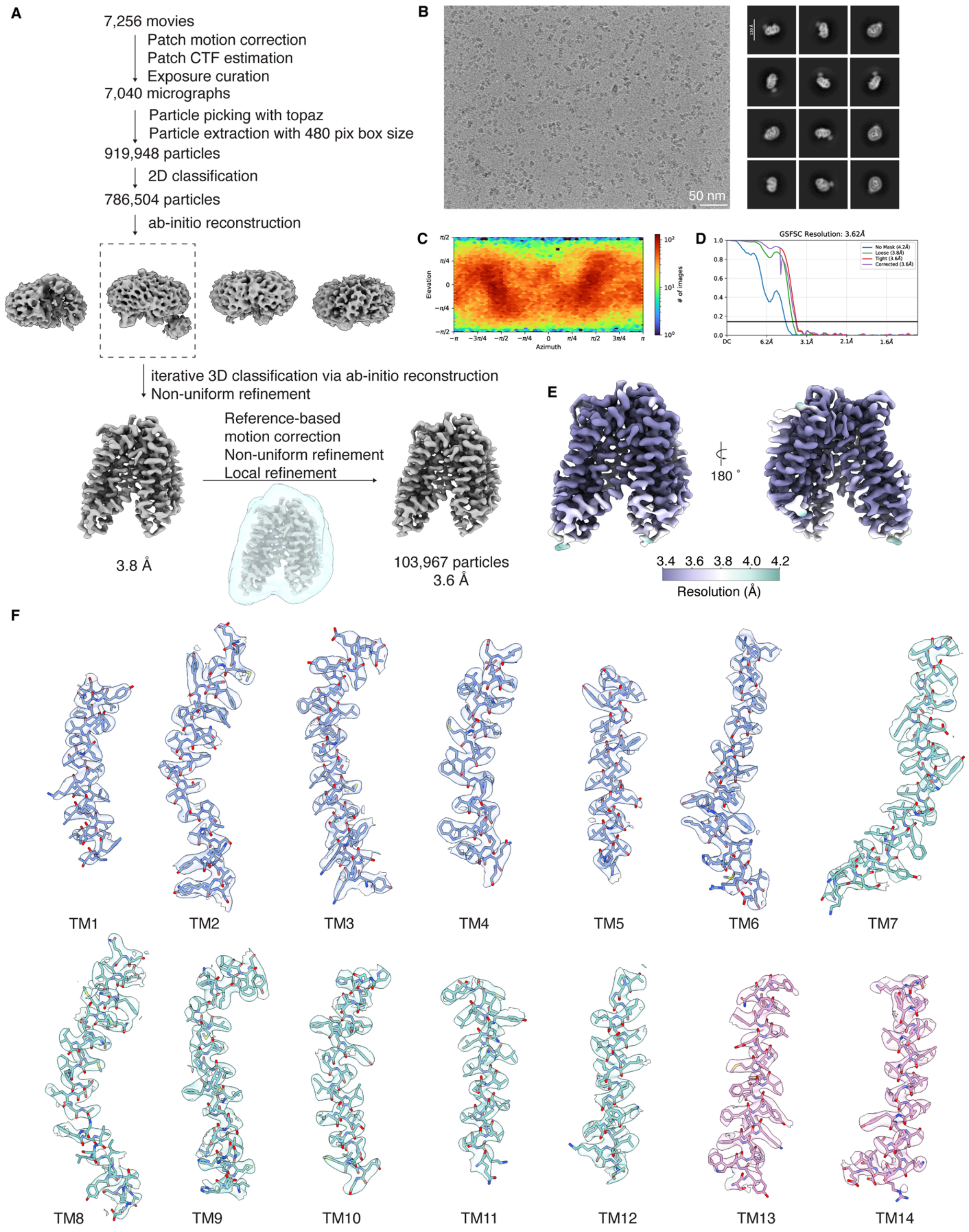

A
7,256 movies
Patch motion correction
Patch CTF estimation
Exposure curation
7,040 micrographs
Particle picking with topaz
Particle extraction with 480 pix box size
919,948 particles
2D classification
786,504 particles
ab-initio reconstruction
iterative 3D classification via ab-initio reconstruction
Non-uniform refinement
Reference-based
motion correction
Non-uniform refinement
Local refinement
3.8 Å
103,967 particles
3.6 Å
B
50 nm
C
Elevation
Azimuth
### of images
D
GSFSC Resolution: 3.62Å
No Mask (4.2Å)
Loose (3.8Å)
Tight (3.6Å)
Corrected (3.6Å)
DC
6.2Å
3.1Å
2.1Å
1.6Å
E
180 °
3.4 3.6 3.8 4.0 4.2
Resolution (Å)
F
TM1
TM2
TM3
TM4
TM5
TM6
TM7
TM8
TM9
TM10
TM11
TM12
TM13
TM14

**Fig. S7. Cryo-EM structure determination of *Sa*MurJ$_{ALFA}$ R176A mutant.** (**A**) Summary of cryo-EM data processing workflow to obtain the structure of *Sa*MurJ R176A mutant. (**B**) Representative micrograph (left) and 2D class averages (right). (**C**) Angular distribution of particles of the final 3D reconstruction. (**D**) Fourier shell correlation curves between two half maps. (**E**) Local resolution map of the final cryo-EM map colored by resolution from 3.4 Å (purple) to 4.2 Å (green). (**F**) Cryo-EM density of each TM helix of *Sa*MurJ R176A mutant. Map contour level = 0.08 in ChimeraX **(4)**.

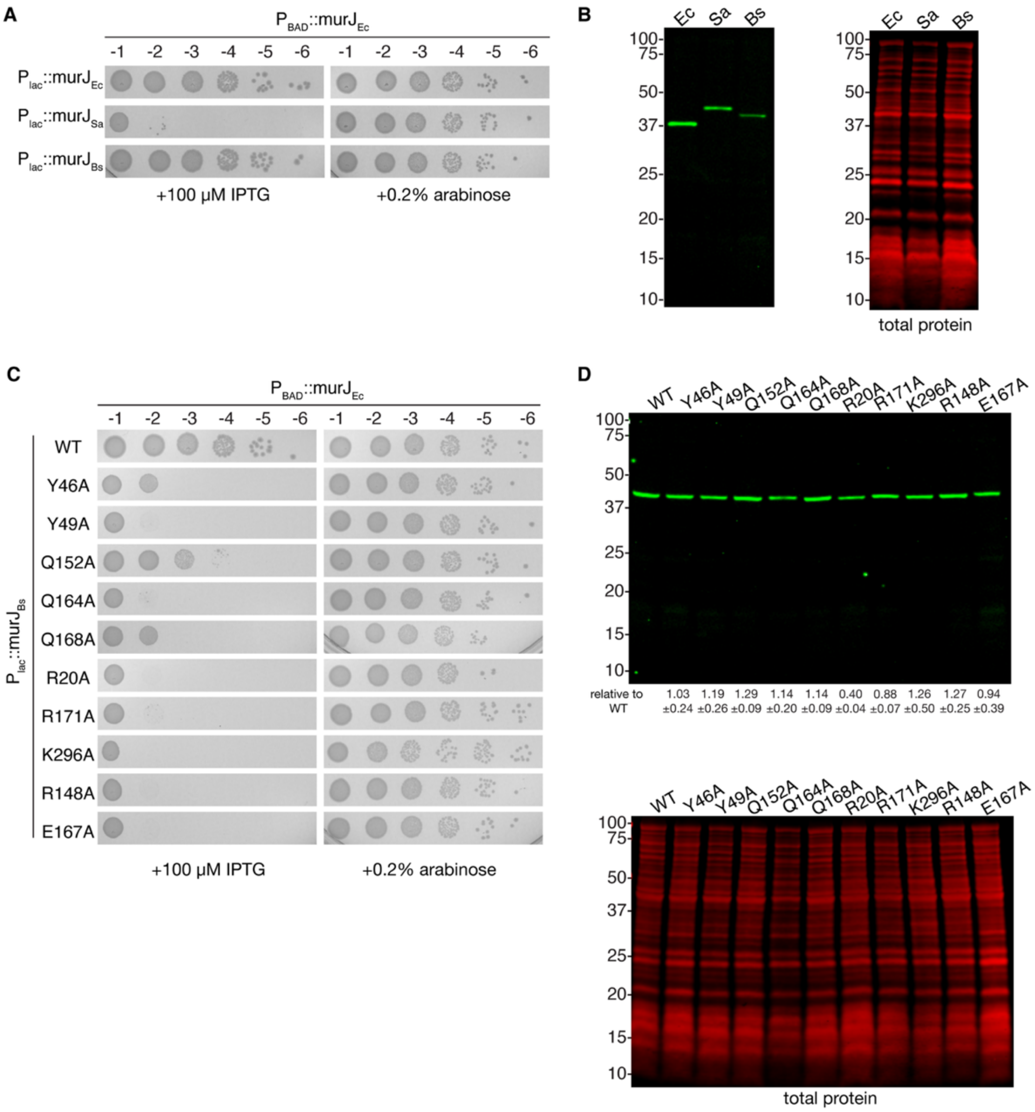


**Fig. S8. Complementation by MurJ homologs and variants.** (**A**) Complementation by MurJ homologs in a MurJ-depletion *E. coli* strain **(5)**. The chromosomal $murJ_{Ec}$ is controlled by an arabinose-inducible promoter. MurJ homologs are expressed from an IPTG-inducible $P_{lac}$ promoter. (**B**) Representative western blots of the membrane fractions from *E. coli* cells expressing different MurJ homologs. Molecular weight markers are indicated for all western blots. (**C**) Complementation by *Bs*MurJ mutants. (**D**) Representative western blots of the membrane fractions from *E. coli* cells expressing *Bs*MurJ mutants. Values below each lane indicate the protein level normalized to total protein staining relative to wild-type, shown as mean ± s.d. from three biological replicates.

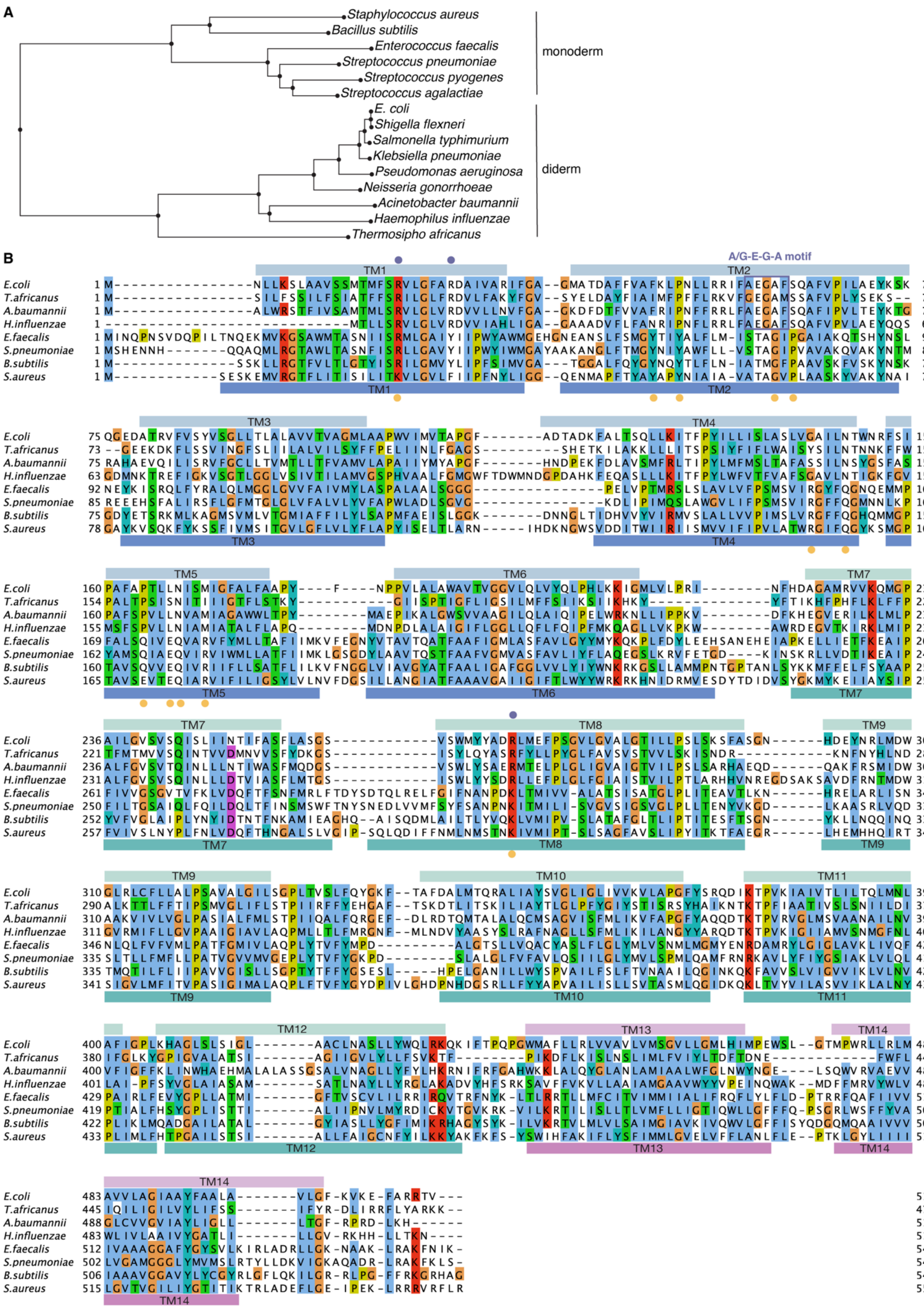
A
Staphylococcus aureus
Bacillus subtilis
Enterococcus faecalis
Streptococcus pneumoniae
Streptococcus pyogenes
Streptococcus agalactiae
E. coli
Shigella flexneri
Salmonella typhimurium
Klebsiella pneumoniae
Pseudomonas aeruginosa
Neisseria gonorrhoeae
Acinetobacter baumannii
Haemophilus influenzae
Thermosipho africanus
monoderm
diderm
B
A/G-E-G-A motif
TM1
TM2
TM3
TM4
TM5
TM6
TM7
TM8
TM9
TM10
TM11
TM12
TM13
TM14
E.coli
T.africanus
A.baumannii
H.influenzae
E.faecalis
S.pneumoniae
B.subtilis
S.aureus

**Figure S9. Sequence alignment of representative MurJ homologs from priority bacterial pathogens. (A)** Phylogenetic tree of representative monoderm and diderm MurJ homologs from the WHO Bacterial Priority Pathogens List 2024 **(6)**. *Thermosipho africanus* is included for completeness. **(B)** Structure-based multiple sequence alignment using Promals3D **(7)** with ClustalW coloring for residues. Species were selected from the phylogenetic tree in (A). MurJ homologs are (Uniprot IDs in brackets): *Escherichia coli* (P0AF16), *Acinetobacter baumannii* (D0CEW3), *Haemophilus influenzae* (P44958), *Thermosipho africanus* (B7IE18), *Staphylococcus aureus* (Q2FXH6), *Enterococcus faecalis* (Q838A3), *Streptococcus pneumoniae* (Q8DP32), and *Bacillus subtilis* (O34674). Secondary structures are shown above the sequence for *Ec*MurJ and below the sequence for *Sa*MurJ colored as in Fig. 1. Functionally important residues are highlighted by circles (purple: *Ec*MurJ, yellow: *Sa*MurJ), including conserved residues implicated in lipid II coordination and conformational rearrangements.

#### Table S1. Cryo-EM data collection, refinement, and validation statistics

| | *Sa*MurJ outward (EMD-76175) (PDB 11XX) | *Ec*MurJ outward (EMD-76176) (PDB 11XY) | *Sa*MurJ inward (EMD-76177) (PDB 11XZ) | *Sa*MurJ R176A (EMD-76178) (PDB 11YA) |
|---|---|---|---|---|
| **Data collection and processing** | | | | |
| Microscope | FEI Titan Krios | FEI Titan Krios | FEI Titan Krios | FEI Titan Krios |
| Magnification | 130,000 | 130,000 | 130,000 | 130,000 |
| Voltage (kV) | 300 | 300 | 300 | 300 |
| Electron exposure (e–/Å$^2$) | 70 | 70 | 70 | 70 |
| Defocus range (μm) | 1.0 to 3.0 | 1.0 to 3.0 | 1.0 to 3.0 | 1.0 to 3.0 |
| Pixel size (Å) | 0.649 | 0.649 | 0.649 | 0.649 |
| Symmetry imposed | C1 | C1 | C1 | C1 |
| Initial particle images (no.) | 508,778 | 833,795 | 361,173 | 919,948 |
| Final particle images (no.) | 57,586 | 72,314 | 92,846 | 103.967 |
| Map resolution (Å) | 3.6 | 3.9 | 3.5 | 3.6 |
| FSC threshold | 0.143 | 0.143 | 0.143 | 0.143 |
| Map sharpening *B* factor (Å$^2$) | −127.1 | −160.4 | −122.3 | −142.9 |
| **Refinement** | | | | |
| Initial model used (PDB code) | SaMurJ AF model | 9NU4 | SaMurJ AF model | SaMurJ AF model |
| Model resolution (Å) | 3.8 | 4.0 | 3.7 | 3.8 |
| FSC threshold | 0.5 | 0.5 | 0.5 | 0.5 |
| Model composition | | | | |
| Non-hydrogen atoms | 4382 | 3792 | 4327 | 4303 |
| Protein residues | 553 | 499 | 546 | 544 |
| Ligands | 0 | 0 | 0 | 0 |
| *B* factors (Å$^2$) | | | | |
| Protein | 78.68 | 62.41 | 54.32 | 64.08 |
| Ligand | --- | --- | --- | --- |
| R.m.s. deviations | | | | |
| Bond lengths (Å) | 0.006 | 0.006 | 0.008 | 0.005 |
| Bond angles (°) | 0.697 | 0.819 | 0.852 | 0.699 |
| Validation | | | | |
| MolProbity score | 1.38 | 1.43 | 1.40 | 1.28 |
| Clashscore | 6.92 | 7.81 | 7.24 | 5.23 |
| Poor rotamers (%) | 0 | 0 | 0 | 0 |
| Ramachandran plot | | | | |
| Favored (%) | 98.19 | 98.59 | 98.53 | 98.15 |
| Allowed (%) | 1.81 | 1.41 | 1.47 | 1.85 |
| Disallowed (%) | 0 | 0 | 0 | 0 |

**SI References**

1. A. C. Y. Kuk, A. Hao, Z. Guan, S.-Y. Lee, Visualizing conformation transitions of the Lipid II flippase MurJ. *Nat Commun* **10**, 1736 (2019).
2. S. Zheng, *et al.*, Structure and mutagenic analysis of the lipid II flippase MurJ from Escherichia coli. *Proceedings of the National Academy of Sciences* **115**, 201802192 (2018).
3. H. Kohga, *et al.*, Crystal structure of the lipid flippase MurJ in a “squeezed” form distinct from its inward- and outward-facing forms. *Structure* (2022). https://doi.org/10.1016/j.str.2022.05.008.
4. T. D. Goddard, *et al.*, UCSF ChimeraX: Meeting modern challenges in visualization and analysis. *Protein Science* **27**, 14–25 (2018).
5. L.-T. Sham, *et al.*, MurJ is the flippase of lipid-linked precursors for peptidoglycan biogenesis. *Science* **345**, 220–222 (2014).
6. *WHO Bacterial Priority Pathogens List 2024: Bacterial Pathogens of Public Health Importance, to Guide Research, Development, and Strategies to Prevent and Control Antimicrobial Resistance*, 1st ed (World Health Organization, 2024).
7. J. Pei, B.-H. Kim, N. V. Grishin, PROMALS3D: a tool for multiple protein sequence and structure alignments. *Nucleic Acids Research* **36**, 2295–2300 (2008).
